# Chilling out or heating up? Thermal plasticity of seminal fluid proteins in *Drosophila melanogaster*

**DOI:** 10.1101/2025.09.26.678772

**Authors:** Claudia Londoño-Nieto, Roberto García-Roa, Stuart Wigby, Irem Sepil, Pau Carazo

## Abstract

Seminal fluid proteins (SFPs) play crucial roles in reproduction by shaping female post-mating physiology and behaviour, male sperm competition, and sexual conflict between the sexes. However, we mostly ignore how abiotic ecological factors regulate SFP expression and transfer. Here, we use quantitative proteomics in adult flies from a wild *Drosophila melanogaster* population to test how exposure to natural temperature variation (20°C, 24°C and 28°C) across two-time scales (48h and 13d), and under either no or high sperm competition risk, interact to shape SFP production and transfer. We show that both production and transfer of SFPs was reduced after long-term (13d) exposure to low (20°C) and high (28°C) temperatures, including in key proteins (sex peptide and ovulin networks) mediating female post-copulatory responses, sperm competition and sexual conflict. Our results show that natural temperature fluctuations can have a strong effect on SFP production and transfer, and thus on post-copulatory sexual selection dynamics.

## Introduction

The last couple of decades have witnessed a revolution in our understanding of the complexity and function of male ejaculates, much of which has been driven by an increasing realization of the importance of non-sperm components (Gillott, 2003; Perry et al., 2013; Wedell et al., 2002). Sperm is not transferred in isolation within the ejaculate, but alongside a wide variety of molecules, including proteins (seminal fluid proteins: SFPs) that are crucial for reproduction (Hopkins et al., 2017). Produced in the male reproductive organs and transferred to the female during mating, SFPs play essential roles in fertility, sperm storage and usage, and strongly impact the fitness of both sexes by affecting female physiology, behaviour, immunity, and life history, as well as biasing the outcome of sperm competition (Avila et al., 2011; Gillott, 2003; Hopkins & Perry, 2022; Perry et al., 2013; Poiani, 2006). SFPs are thus fundamental to male and female reproductive success and are subject to strong natural and sexual selection (Perry et al., 2013; Sirot et al., 2015). For example, SFPs can alter female physiology and behaviour in ways that are costly to the female, yet beneficial to the male, making them a case study in sexual conflict (Sirot et al., 2015).

Sexual conflict arises because, in sexually reproducing organisms, anisogamy often leads to distinct sex roles (Janicke et al., 2016), which in turn result in divergent reproductive interests between the sexes (Arnqvist & Rowe, 2005; Pizzari & Snook, 2003). Sexual conflict is particularly intense in polygynous species, where it frequently results in male adaptations that increase male fitness at the expense of harming females (i.e. male harm; <u>Arnqvist & Rowe, 2005)</u>. Male harming of females is taxonomically widespread and can significantly depress net female productivity, with potentially substantial demographic consequences for populations (Flintham et al., 2023; Gómez-Llano et al., 2024; Martinez-Ruiz & Knell, 2017). An important avenue of male harm are male adaptations driven by strong sperm competition, such as traumatic insemination and “harmful” SFPs (Mueller et al., 2007; Perry et al., 2013; Reinhardt et al., 2015). Work on *Drosophila melanogaster* has been at the forefront of this field (Wigby et al., 2020), providing compelling evidence that individual SFPs are sufficient to alter female post-mating physiology and behaviour (Hopkins & Perry, 2022). For example, SFPs such as sex peptide (SP) and ovulin regulate key female post-mating responses. SP stimulates oogenesis, reduces female receptivity to remating, and facilitates sperm storage and release (Chapman, 2001; Chapman et al., 2003; Liu & Kubli, 2003; Soller et al., 1999), while ovulin increases the rate of release of existing eggs (ovulation) by the ovaries. Furthermore, other proteins in the SP/ovulin network: *antr, aqrs, intr, lectin-46Ca, lectin-46Cb, Sems, CG17575* and *CG9997*, contribute to the long-term persistence of reduced female receptivity to remating, effective sperm release, and fecundity stimulation (Findlay et al., 2014; Ravi Ram & Wolfner, 2007), while *Semp1* is necessary to process ovulin in mated females (Ravi Ram et al., 2006). Together, SP, ovulin and their associated network proteins increase male reproductive success, and SP and its network proteins are believed to be under sexual conflict, as they increase male fitness but can substantially reduce female lifespan and long-term reproductive output (Wigby & Chapman, 2005).

There is increasing evidence that SFPs production and transfer is highly plastic. A male’s state, socio-sexual environment and condition can affect both the composition and allocation strategies of his seminal fluid proteome (Wigby et al., 2009). *Drosophila* males, for instance, can tailor the quantity of SFPs they transfer in response to perceived sperm competition risk (Hopkins et al., 2019), and may even exploit SFPs from rival ejaculates to maximize their own reproductive success (Sirot et al., 2011). There is also evidence that SFPs’ post-mating effects on females can be highly plastic themselves (Hopkins & Perry, 2022). However, our understanding of how abiotic ecological factors influence the production and transfer of SFPs, and how these interact with sexual selection and conflict, remain limited. This gap persists despite mounting evidence suggesting that the environment is a crucial determinant of both fertility and competitive fertilisation success (Sales et al., 2018; Vasudeva et al., 2014; Yun et al., 2017). In particular, responses of SFPs to environmental variables are prime candidates for mediating these effects. For example, studies show that the larval environment can shape SFP allocation patterns later in adult life (Von Hellfeld et al., 2025; Wigby et al., 2016).

Temperature is a particularly salient ecological factor in this respect, due to its potentially far-reaching effects on sexual reproduction and sexual selection/conflict processes (Dougherty et al., 2024; García-Roa et al., 2020). In nature, temperature is subject to substantial spatiotemporal variability at different scales (e.g. circadian, seasonal), potentially imposing strong selective pressures on sperm and ejaculate traits related to sexual selection and reproductive performance, especially in ectothermic species. For instance, high temperatures can impair gonad development, reduce sperm production, viability and function, (Peña et al., 2019; Vasudeva et al., 2014; Wang & Gunderson, 2022), and compromise protein stability and function (Berger et al., 2021), potentially modifying the efficacy of SFPs (Canal Domenech & Fricke, 2022). However, thermal stress may also impair female physiological defences against male manipulation, suggesting that temperature could influence both male persistence and female resistance mechanisms. Thus, the expression and consequences of sexual conflict could change not only quantitatively but qualitatively across thermal environments. If thermal variation compromises SFP production and transfer, it may reduce male influence over female post-mating responses, altering the balance between pre- and post-copulatory sexual selection. Recent evidence in *D. melanogaster* shows that even a small temperature increase, well within the wild females’ optimal reproductive range under no sexual conflict (i.e. from 24°C to 28°C), is sufficient to reduce both the impact of male ejaculates on female receptivity and overall male harm to females under high sexual conflict (Londoño-Nieto et al., 2023). However, it remains unclear whether such reductions in ejaculate-mediated female responses arise because males produce or transfer different abundances of SFPs, or because temperature alters the functional effects of these proteins, both within the male and after transfer. Furthermore, simultaneously coping with thermal variation and variation in SCR may generate physiological trade-offs for males. Therefore, understanding how temperature influences SFP dynamics across different socio-sexual contexts is crucial to understand how environmental variation shapes male ejaculate composition and sexual selection dynamics, and may also offer important insight into the thermal limits of fertility (Dougherty et al., 2024).

Here, we used *D. melanogaster* flies to investigate how natural temperature variation, within the optimal reproductive range for a wild population, affects male production and transfer of SFPs, and whether these effects change with perceived sperm competition risk (hereafter SCR). We assessed these effects by exposing adult males to a factorial combination of temperature (i.e. 20°C, 24°C and 28°C) and SCR treatments (i.e. no-SCR vs. high-SCR) across two different time scales (experiment 1: short-term exposure-48h; experiment 2: long-term exposure-13d). We predicted that the seminal fluid proteome would be thermally plastic, with 28°C impacting the production and/or transfer of SFP, particularly after long-term exposure (Berger et al., 2021; Zheng et al., 2024). We also predicted that production/transfer of SFPs under sexual conflict (i.e. the sex peptide, ovulin and their network proteins) would be generally reduced at 28°C and 20°C after long-term exposure, based on the fact that male harm is reduced after long-term exposure to 20°C and 28°C relative to 24°C in *Drosophila melanogaster* (García-Roa et al., 2019; Londoño-Nieto et al., 2023). More specifically, we predicted that long-term exposure to 28°C would reduce the abundance of functionally of important SFPs involved in mediating female receptivity (i.e. *SP*, *antr, aqrs, intr, lectin-46Ca, lectin-46Cb, Sems, CG17575* and *CG9997*), given the reduced impact of male ejaculates on female receptivity after long-term exposure to 28°C (Londoño-Nieto et al., 2023).

## Materials and Methods

### Stock Maintenance

We carried out all experiments using individuals from our field-collected population, “Vegalibre”, established in October 2018 from three wineries in Requena (Spain), and re-sampled annually to maintain natural genetic variation (Londoño-Nieto et al., 2023). We keep Vegalibre in the laboratory with overlapping generations at an average temperature of 24°C with daily pre-programmed fluctuations (±4°C) mimicking natural daily temperature conditions during the reproductively active season, at ∼60% humidity and on a 12:12 hr light:dark cycle. We used maize-malt medium (see Londoño-Nieto et al., 2023) as a food source throughout maintenance and experiments. To collect experimental flies, we introduced yeasted grape juice agar plates into stock populations to induce female oviposition. We then collected eggs and placed them in bottles containing ∼75 ml of medium to be incubated at 24 ± 4°C at a controlled density (Clancy & Kennington, 2001). We collected virgin flies within 6 hr of eclosion, under ice anesthesia, and then sexed and kept them in vials with food until their use (see below). All females used in our experiments were 4 days old, while males were 4 days old for experiment 1 and 16 days old for experiment 2, at the time when samples were collected.

### Proteomics experiments

#### Experimental overview and detailed methods

To study the effect of temperature on the production and transfer of SFPs across different reproductive contexts, we exposed adult male flies to either no or high precopulatory levels of SCR (Hopkins et al., 2019), across three stable temperature treatments of 20°C, 24°C, and 28°C, for two different treatment durations/scales.

In experiment 1, we simulated short-term temperature effects (intragenerational effects) on the production and transfer of SFPs. Upon eclosion, we randomly placed virgin males either individually (no-SCR) or in a group of 8 males (high-SCR) in plastic vials with food. Next, we randomly divided vials into three groups that we allocated to one of the three stable temperature treatments (20°C, 24°C, and 28°C) for 48-hours (Fig. S1).

In experiment 2, we examined longer-term temperature effects on the production and transfer of newly synthesized SFPs that would be typical of seasonal (intra- and inter-generational) effects. We took 2 days-old males and first depleted their ejaculates by housing them individually with four virgin females for 24 hours, since *D. melanogaster* males can become sperm and SFP depleted when mated sequentially over a short period (Sirot et al., 2009). Then, we placed the 3 days-old depleted males for 13 days in the same temperature-SCR treatments of the experiment 1 (Fig. S2). Because males take at least 3 days to fully replenish their SFPs after depletion (Sirot et al., 2009), this design ensured that males synthesized new SFPs under our temperature treatments.

#### Quantitative proteomics

Immediately after treating males, we explored SFP production patterns at a common garden temperature of 24°C, by measuring the abundance of SFPs in virgin (experiment 1) or recently unmated (for 13d, experiment 2) males across all temperature-SCR treatment combinations. To explore SFP transfer patterns, we compared the abundance of SFPs in virgin/recently unmated males vs. recently mated males across the same treatment combinations (Hopkins et al., 2019; Sepil et al., 2019, 2020).

Four days before sample collection, we collected females as virgins and held them in same sex groups of 15-20 flies at 24±4°C. On the day of sample collection, we isolated 240 females in vials with food, after which we immediately introduced one focal male either into a vial containing a female or into an empty vial to be retained as a virgin. We flash froze recently mated males in liquid nitrogen 25 min after the start of mating, freezing a virgin/recently unmated male from the same temperature-SCR treatment combination at the same time (Figs. S1-2). Thus, the distribution of freezing times among virgin/recently unmated and recently mated males was equivalent. Freezing males at 25 min after the start of mating ensures a complete mating, where mating typically last ∼20min (see Hopkins et al., 2019; Sepil et al., 2019). We flash froze 40 males per temperature-SCR treatment combination (20 virgin/recently unmated and 20 recently mated). To obtain three independent biological replicates, we repeated this procedure over three different cohorts from the same stock population on different days (i.e. blocks) for each experiment, with all SCR and temperature treatments equally represented within each block.

We stored all frozen samples at -80°C until dissection, for which we thawed flash frozen males and dissected their accessory glands, severing the ejaculatory duct and removing the seminal vesicles and testes on ice in phosphate-buffered saline (PBS) under a binocular scope (Hopkins et al., 2019). Each biological replicate (i.e. sample) consisted of a pool of 20 reproductive glands in 25-µL PBS buffer. Samples were analysed with label-free quantitative proteomics, using the sample preparation and quantification protocol (SWATH-MS; Gillet et al., 2012) at the SCISIE proteomics service at the University of Valencia. Samples were centrifuged and protein pellets resuspended in Laemmli buffer. Proteins were separated by SDS-PAGE, full gel separation was performed for spectral library construction, while a short gel run was used to clean and concentrate individual samples, followed by in-gel trypsin digestion as described by <u>Shevchenko et al. (1996)</u>. Peptides were analysed using a nanoESI-qQTOF-6600+ TripleTOF mass spectrometer (AB SCIEX) coupled to a nanoflow LC system. Protein identification and quantification were carried out using ProteinPilot and PeakView software with a 1% false discovery rate threshold. Full methodological details are available in SI. We analysed six samples per temperature treatment-SCR combination (three from virgin or recently unmated males and three from recently mated males). In total, we had 72 samples (36 from experiment 1 and 36 from experiment 2) from 1440 males.

### Data analysis

We conducted all proteomics analysis on normalized abundances. We normalized protein areas calculated by dividing each protein abundance by the total sum of the abundance of all quantified proteins. We excluded proteins with fewer than 2 unique peptides to restrict analysis to proteins with high-confidence quantification (Carr et al., 2004). We identified SFPs based on the high-confidence SFPs reference list from Wigby et al. (2020). Given the high dimensionality of the dataset and the limited sample size, we adopted an analytical strategy combining Elastic net regularization and targeted, hypothesis-driven Bayesian multivariate analysis. Classical linear models were deemed unsuitable for the full dataset due to the high risk of overparameterization, and the resulting loss of statistical power, a constraint inherent to proteomics datasets of this type, where the number of variables routinely exceeds the number of biological replicates (i.e. assuming a 1.5-fold change in protein abundance and using the dataset with the lowest residual variance across treatments, power was below 0.1). We therefore implemented an Elastic net penalized multinomial regression using *glmnet* (Friedman et al., 2010) and *caret* (Kuhn, 2008) packages in RStudio. Elastic net is a hybrid regularization method that combines Lasso and Ridge penalties to overcome severe multicollinearity (Zou & Hastie, 2005), a classic feature of highly co-expressed proteomic datasets. It is particularly suited for situations where the number of predictors far exceeds the number of observations, as it performs both regularization and variable selection. Consequently, the method sets weakly associated or irrelevant predictors to exactly zero, while preserving coordinated groups of highly correlated variables with stronger effects via non-zero coefficients. Model tuning parameters: α, which balances the contribution of Lasso (α=1) and Ridge (α=0), and λ, which determines the overall penalty strength, were optimized using repeated cross-validation (Zou & Hastie, 2005). Under this framework, proteins with non-zero coefficients were interpreted as the multivariate signature driving variation across temperature treatments.

To avoid overparameterization and maximize statistical power, we broke down analyses for both experiments using the following scheme. To analyse how temperature affects SFP production patterns (i.e. the abundance of SFPs in the accessory glands of virgin –in experiment 1– or recently unmated –in experiment 2– males), we generated two datasets (one per SCR level), including all virgin (experiment 1) or recently unmated (experiment 2) male samples across all temperature treatments. To analyse how temperature influences SFP transfer, we first inferred the relative abundances of transferred SFPs by dividing the abundance of each SFP in virgin or recently unmated males by its abundance in recently mated males within the same temperature treatment, SCR level, and replicate. Based on these ratios, we then generated two datasets (one per SCR level) including SFP transfer estimates across all temperature treatments. Note, however, that we also analysed absolute transfer of SFPs (calculated as the difference between virgin or recently unmated males and recently mated males), yielding qualitatively similar results (see SI). To visualize the abundance of SFPs produced and transferred, we took the average across the three biological replicates for each SFP within each temperature treatment and SCR level, and then generated heatmaps using a Euclidean distance metric and plotted with the aheatmap function from the *NMF* package (Gaujoux & Seoighe, 2010) in RStudio (version R 4.2.2; R Core Team, 2022). We also constructed population mean reaction norms, allowing for clearer visualization of directional trends, plastic responses, and potential non-linear expression patterns. Finally, we performed a targeted analysis strictly on a subset of proteins including the SP and Ovulin, along with their associated protein networks (*antr, aqrs, intr, lectin-46Ca, lectin-46Cb, Sems, CG17575, CG9997* and *Semp1)* which have well-established roles in female post-mating responses (Findlay et al., 2014; Ravi Ram & Wolfner, 2007; Ravi Ram et al., 2006; Liu & Kubli, 2003). We fitted separate multivariate Bayesian linear models for production and transfer data from each of four assays, using the *brms* package (Bürkner, 2017) interfaced to Stan in RStudio. In each model, the log(1+x)-transformed abundance of each SFP was specified as a separate response variable within a joint multivariate framework, allowing residual correlations among proteins to be estimated simultaneously. Temperature (20°C, 24°C, and 28°C) was treated as a categorical fixed effect, with 24°C as the reference level, yielding two contrast parameters per protein (20°C vs. 24°C and 28°C vs. 24°C). We used weakly informative priors (i.e. priors that constrain estimates to plausible ranges without strongly favouring any particular value, allowing the data to drive inference) on regression coefficients and intercepts, default half-Student-t priors on residual standard deviations and an LKJ (1) prior on the residual correlation matrix (i.e. no prior belief about whether proteins are positively correlated, negatively correlated, or uncorrelated with each other). Models were fitted with four chains of 3,000 iterations each (1,500 warmup), and a target acceptance rate of adaptdelta = 0.99, except for one model (Exp 1 high-SCR production) that was refitted with 6,000 iterations and adaptdelta = 0.995 to resolve mixing issues encountered in the initial run. Convergence was confirmed using the potential scale reduction factor (R^ < 1.01) and effective sample sizes. Temperature effects are reported as posterior means with 95% credible intervals (CrI) and posterior probabilities of direction [P(β > 0)]; effects with 95% CrI excluding zero were interpreted as strong evidence of a temperature-driven change in SFP production or transfer.

## Results

We found a total of 1492 proteins, of which we retained 1328 with 2 or more unique peptides. Among these, we focused on the 145 previously identified as SFPs.

### Temperature effects on the overall plasticity of SFP production and transfer

Our results show that SFP production and transfer are highly plastic in response to temperature variation across both treatment durations, but with important differences across experiments. In males treated for 48-hours (short-term -experiment 1), Elastic net regression identified 74 (under no-SCR) and 3 (under high-SCR) SFPs as predictors contributing most strongly to temperature variation in protein production (see Tables S2 and S3), and 79 (under no-SCR) and 6 (under high-SCR) in the case of SFP transfer (Tables S4 and S5). Heatmaps revealed that both production and transfer of SFPs changed with temperature, with clearer effects under no-SCR. Population mean reaction norms supported these findings (Fig. 1), suggesting that SFP production and transfer were more strongly affected by temperature in males exposed to no-SCR than in those under high-SCR. Both the number of SFPs showing temperature-mediated changes in abundance and the magnitude of these changes were markedly reduced under high-SCR (Fig. 1), suggesting that under high-SCR male ejaculates are less thermally plastic after a short-term exposure. In males treated for 13-days (long-term -experiment 2), Elastic net regression selected 39 (under no-SCR) and 68 (under high-SCR) SFPs for production (Tables S6 and S7), and 49 (under no-SCR) and 46 (under high-SCR) SFPs for transfer (Tables S8 and S9) as predictors contributing most strongly to differential response to temperature. Heatmap inspection showed clear temperature-driven variation in SFP production and transfer, confirmed by the population mean reaction norms (Fig. 2). Overall, we found clear evidence that temperature effects after long-term exposure were more uniform across SCR conditions, consistently reducing SFP production and transfer at both low (20°C) and hot (28°C) temperatures (Fig. 2). Estimates of SFP transfer derived from absolute abundances produced comparable heatmap patterns, supporting the observed temperature effects (Figs. S3-4).

**Figure 1.**
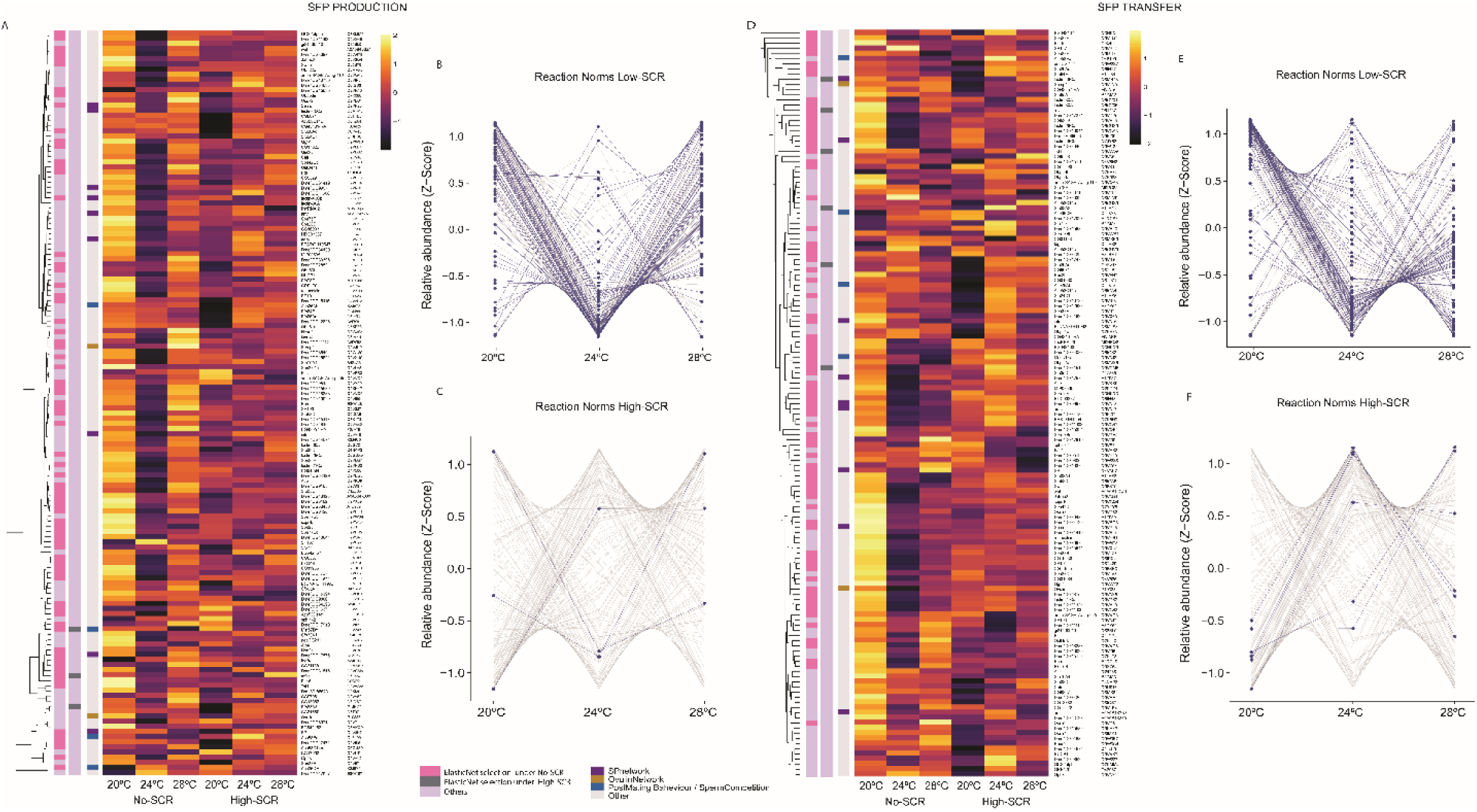
Effect of temperature on SFPs production and transfer (short-term exposure / experiment 1). A) Heatmap showing the abundance of 145 SFPs detected in accessory gland samples. Elastic net regression selected 74 (under no-SCR) and 3 (under High-SCR) out of 145 SFPs as predictors with a strong effect differentiating proteomic responses across temperature treatments. Each cell gives the across-biological replicate mean for that protein, at each temperature treatment. Row annotations indicate SFPs with strong response to temperature in each SCR level and include functional information relating to protein functions as part of the sex peptide or ovulin networks, or other known roles in sperm competition. B) Population mean reaction norms of SFP production under no-SCR treatment and C) under high-SCR treatment across temperature treatments. Faint grey lines represent the global SFP background using standardized abundances. Proteins selected by multivariate Elastic net models are highlighted with blue lines and solid points. D) Heatmap showing the inferred relative abundance of 145 SFPs transferred to females during mating. Elastic net regression selected 79 (no-SCR treatment) and 6 (High-SCR treatment) out of 145 SFPs as predictors with strong differential transfer response to temperature. Each cell shows the mean ratio of protein abundance in virgin males relative to recently mated males, across biological replicates, for each temperature treatment, reflecting the relative loss of protein upon mating. Heatmaps were generated from log2-transformed protein abundances for better visualization. E) Population mean reaction norms of SFP transfer under no-SCR treatment and F) under high-SCR treatment across temperature treatments. Faint grey lines represent the global SFP background using standardized abundances, with Elastic net selected proteins highlighted by blue lines and solid points.

**Figure 2.**
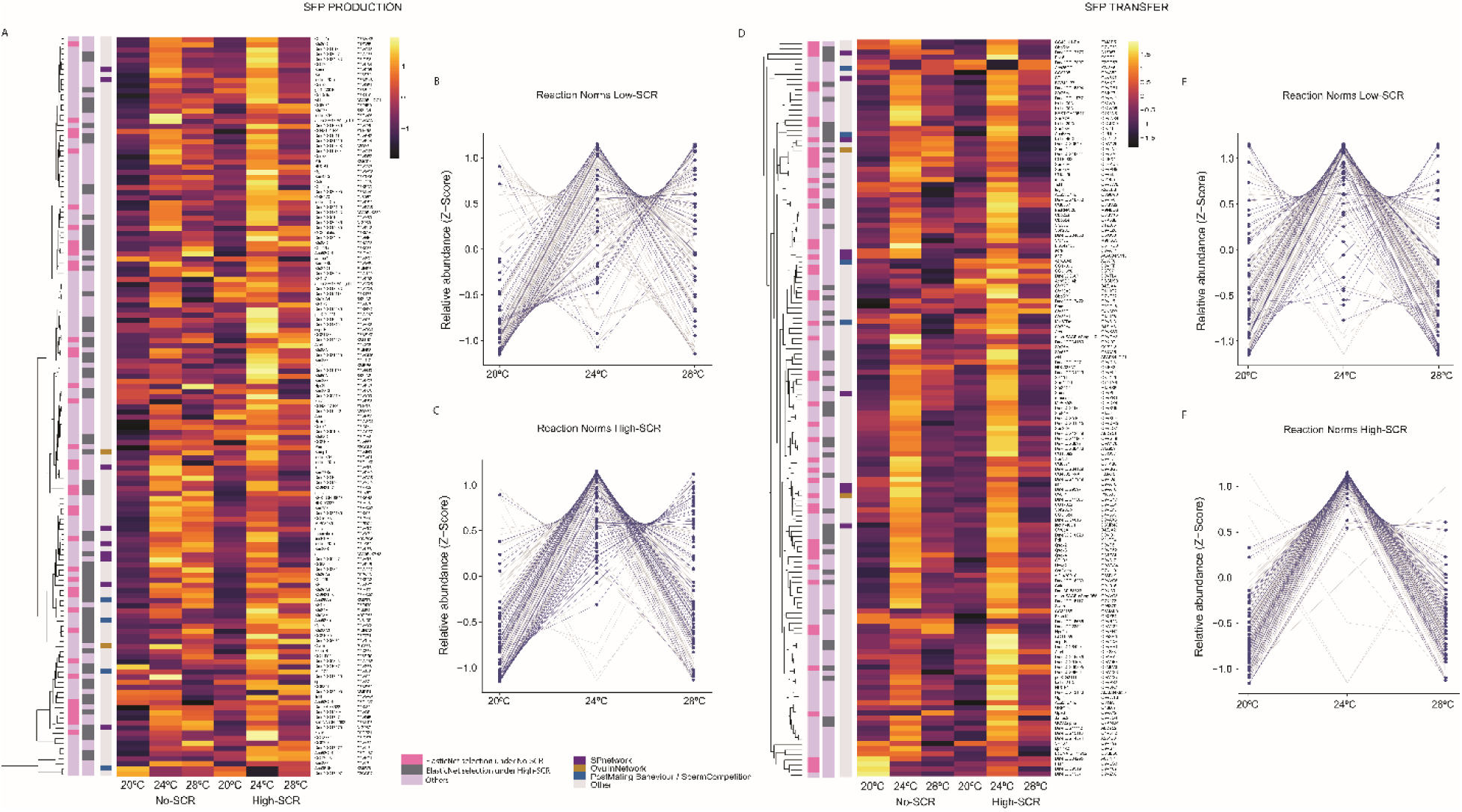
Effect of temperature on SFPs production and transfer (long-term exposure / experiment 2). A) Heatmap showing the abundance of the 145 SFPs detected in the accessory gland samples. Elastic net regression selected 39 (no-SCR treatment) and 68 (high-SCR treatment) out of 145 SFPs as predictors with a strong effect differentiating proteomic responses across temperature treatments. Each cell gives the across-biological replicate mean for that protein, at each temperature treatment. Row annotations indicate SFPs with strong response to temperature in each SCR level and include functional information relating to protein functions as part of the sex peptide or ovulin networks, or other known roles in sperm competition. B) Population mean reaction norms of SFP production under no-SCR treatment and C) under high-SCR treatment across temperature treatments. Faint grey lines represent the global SFP background using standardized abundances. Proteins selected by multivariate Elastic net models are highlighted with blue lines and solid points. D) Heatmap showing the inferred relative abundance of 145 SFPs transferred to females during mating. Elastic net regression selected 49 (no-SCR treatment) and 46 (high-SCR treatment) out of 145 SFPs as predictors with strong differential transfer response to temperature. Each cell shows the mean ratio of protein abundance in unmated males relative to recently mated males, across biological replicates, for each temperature treatment, reflecting the relative loss of protein upon mating. Heatmaps were generated from log2-transformed protein abundances for better visualization. E) Population mean reaction norms of SFP transfer under no-SCR treatment and F) under high-SCR treatment across temperature treatments. Faint grey lines represent the global SFP background using standardized abundances, with Elastic net selected proteins highlighted by blue lines and solid points.

### Temperature effects on the production and transfer of SFPs involved in female post-mating responses

For proteins involved in female post-mating responses, we identified 8 members of the SP network (*antr, aqrs, intr, lectin-46Ca, lectin-46Cb, Sems, CG17575* and *CG9997*) as well as SP itself, ovulin, and its precursor *Semp1.* Our results show that, at 20°C and 28°C, the production and transfer of these proteins, and specifically those involved in the SP network, frequently diverged from that observed at 24°C, with clearer effects after the long-term treatment duration (experiment 2).

In experiment 1 (short-term exposure), SFP production revealed a consistent pattern: protein abundance at 24°C was markedly reduced compared to 20°C and 28°C, but only under no-SCR (Fig. 3A). Among the selected proteins via Elastic net under no-SCR, SP network proteins *Sems, lectin-46Cb* and *CG17575* followed this pattern. In contrast, *SP* decreased and *Semp1* increased at 28°C (Figs. 1A and S5). Under high-SCR we did not identify a consistent pattern. Consistent with differences in production, SFP transfer under no-SCR was generally reduced at 24°C compared to 20°C and 28°C, while we did not find a clear temperature-dependent pattern under high-SCR (Fig. 3B). Moreover, *ovulin, SP* and SP network proteins *lectin-46Cb* and *CG9997* were among those selected by Elastic net as important predictors only under no-SCR (Figs. 1D and S6). Note, however, that the absolute transfer of *SP* was higher at 24°C than at 20° and 28°C (Fig. S3). Under high-SCR, *lectin-46Ca*, a SP-network protein, was the only SFP known to regulate female post-mating physiology selected by the multivariate model, with higher relative and absolute transfer at 28°C compared to lower temperatures (Figs. 1D, S3 and S6).

**Figure 3.**
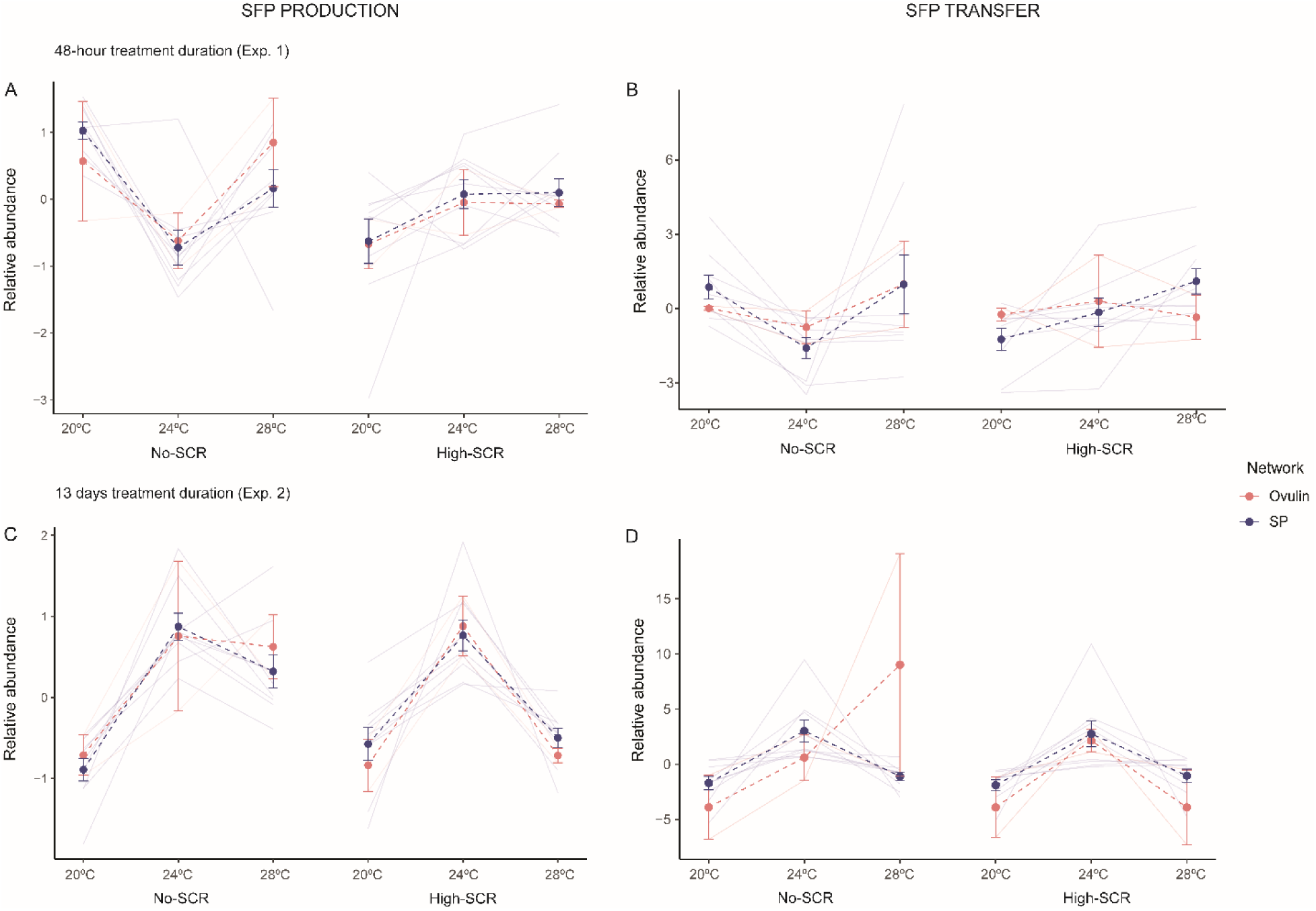
Effect of temperature on production and transfer of SFPs involved in Ovulin and SP networks. A – B) Relative abundance production and transfer profiles of SFPs associated with the SP and ovulin networks across different temperatures and SCR levels for short-term exposure (experiment 1). C – D) Relative abundance production and transfer profiles of SFPs associated with the SP and ovulin networks across different temperatures and SCR levels for long-term exposure (experiment 2). Each faint line represents the mean across the 3 replicates for an individual SFP within each treatment combination. Abundance values were normalized by mean-centering and averaged across replicates. Blue points represent the average (± s.e.) of SFPs in the SP network, across temperature and SCR conditions. Red points represent the average (± s.e.) of SFPs in the ovulin network, across temperature and SCR conditions.

Importantly, Bayesian multivariate analysis of the subset of 11 proteins, including the SP and ovulin, along with their associated protein networks confirmed these dynamics. Under no-SCR, production of most SFPs showed a similar trend of up-regulation at both 20°C and 28°C vs. 24°C, with a clear effect on *Sems* at both temperatures (20°C: mean = 1.43, 95% CrI = 0.47, 2.26; 28°C: mean = 0.97, 95% CrI = 0.01, 1.81) and on *lectin-46Cb* at 20°C only (mean = 1.44; CrI = 0.08, 2.61; Fig. 4A and Table S10A). For transfer, both *CG9997* (mean = 1.10; 95% CrI = 0.27, 1.83) and *lectin-46Cb* (mean = 1.00; 95% CrI = 0.11, 1.81) were robustly up-regulated at 20°C vs. 24°C (Fig. 4B and Table S10B). In contrast, under high-SCR, we found no consistent up/down-regulation in either production or transfer of this protein subset (Fig. S11 and Table S11).

**Figure 4.**
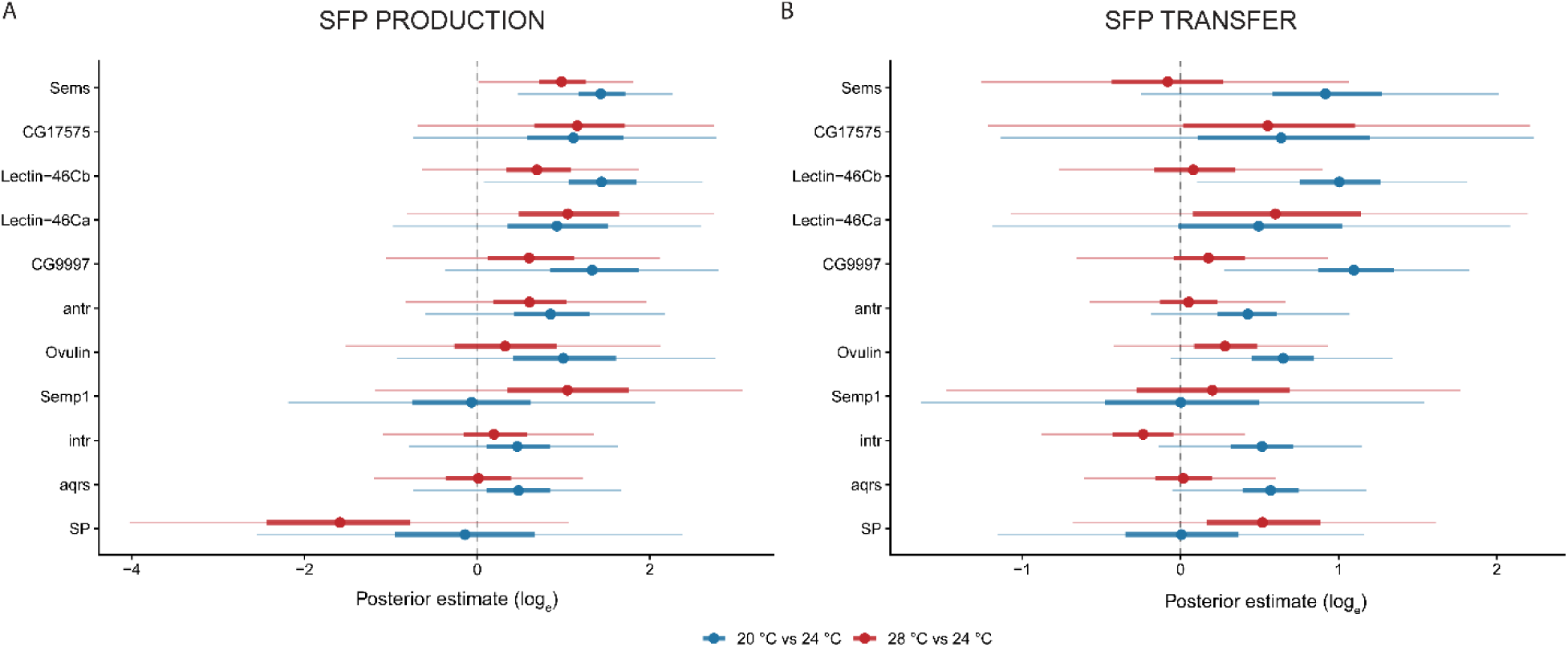
Bayesian posterior estimates of temperature effect on production and transfer of SFPs involved in Ovulin and SP networks (short-term exposure / experiment 1 under no-SCR). log_e_ fold-change estimates of protein abundance induced by low (20°C) and hot (28°C) temperatures relative to the 24°C baseline. A) Posterior estimates for SFP production. B) Posterior estimates for SFP transfer. Blue intervals represent the 20°C vs 24°C contrast, while red intervals represent the 28°C vs 24°C contrast. Central point on each segment denotes the median posterior estimate for a specific protein. Thick and thin horizontal lines represent the 50% and 95% Credible Intervals (CrI), respectively. Vertical dashed line at zero represents no change in abundance compared to the 24°C baseline. Estimates where the 95% CrI strictly excludes zero indicate a highly probable (>95%) robust difference induced by the specific temperature treatment.

In experiment 2 (long-term exposure), SFP production showed a clear trend of increased production at 24°C compared to 20°C and 28°C, regardless of SCR (Fig. 3C). SP and ovulin network proteins (i.e., *Semp1*, *antr*, *SP*, and *CG17575)* were selected as important multivariate predictors only in males under high-SCR (Fig. 2A), showing the general pattern of increased production at 24°C (Figs. 2A and S7). Patterns of SFP transfer mirrored those observed for production. In both SCR conditions, 24°C yielded the highest relative and absolute transfer levels (Figs. 3D and S4). Among the proteins showing temperature-dependent transfer, Elastic net identified *Semp1*, *antr*, *CG17575, lectin-46Cb,* and *Ovulin* in males under no-SCR, and *Semp1*, *CG17575, lectin-46Cb,* and *lectin-46Ca* under high-SCR as those most strongly associated with temperature variation. These proteins exhibited markedly higher transfer at 24°C compared to 20°C and 28°C (Figs. S4 and S8). Despite the general similarity across SCR levels, Elastic net grouped more proteins from the SP and ovulin networks as strongly temperature-sensitive for production under high-SCR, while transfer patterns showed a similar thermal response across SCR levels (Fig. 2). In accordance, Bayesian analysis indicated an overall suppression of both production and transfer across the 11 proteins in the ovulin and SP networks, with the strongest effects on protein transfer under high-SCR. Specifically, under no-SCR, Bayesian estimates identified *intr* as the only protein showing clear evidence of reduced production at 20°C vs 24°C (mean = -0.80, 95% CrI = -1.35, -0.21; Table S12A and Fig. S10A). Under high-SCR, transfer was consistently reduced at both 20°C and 28°C for several proteins. In particular, *CG17575* showed strong down-regulation at 20°C (mean = -2.38; 95% CrI = -3.16, -1.58) and 28°C (mean = -2.22; 95% CrI= -2.99, -1.43).

Similarly, *lectin-46Ca* was reduced at both 20°C (mean = -1.39; 95% CrI = -2.13, -0.62) and 28°C (mean = -1.19; 95% CrI = -1.94, -0.44). *Sems* (20°C mean = -1.16; 95% CrI = -2.32, 0.04; 28°C mean = -1.16; 95% CrI = -2.28, 0.02) and *lectin-46Cb* (20°C mean = -0.99; 95% CrI = -2.04, 0.04; 28°C mean = -2.23; 95% CrI = -2.23, -0.18) showed similar down-regulation trends, although some CrI slightly overlapped zero (Fig. 5B and Table S13B).

**Figure 5.**
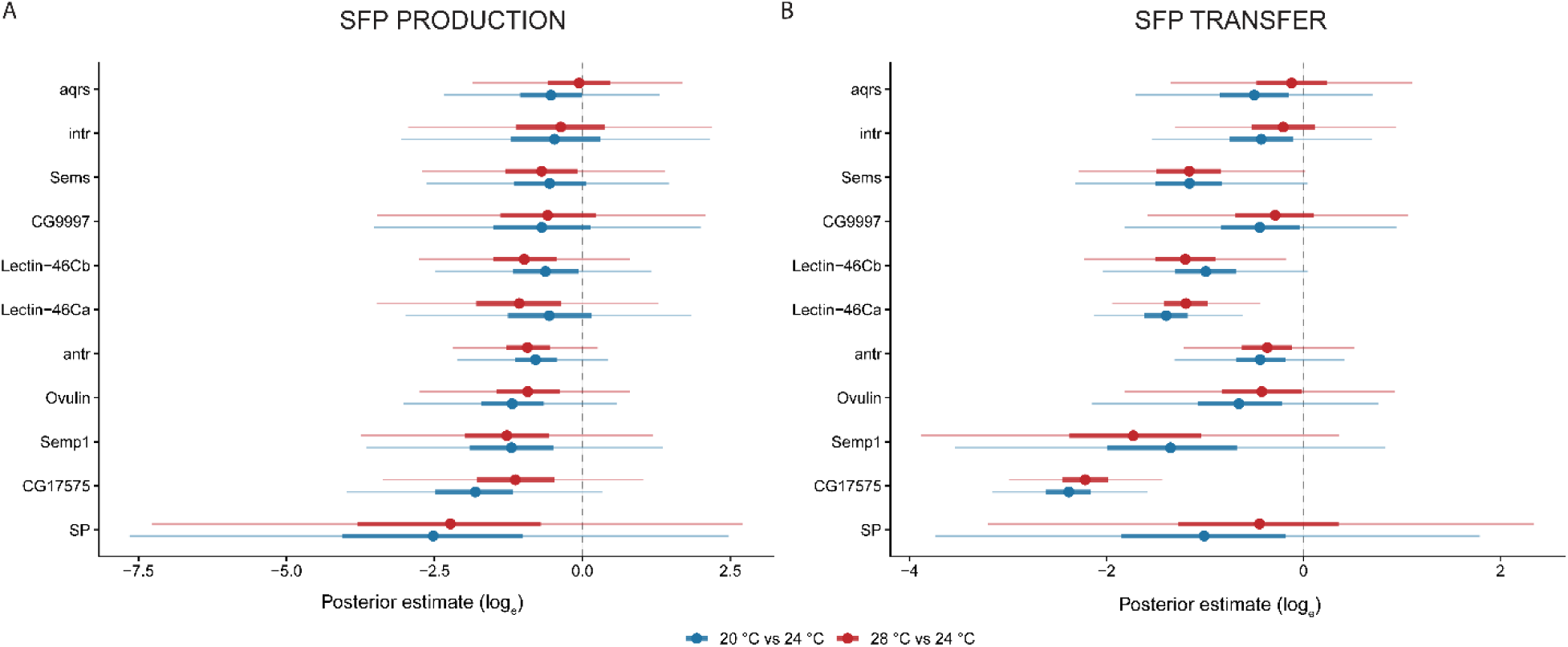
Bayesian posterior estimates of temperature effect on production and transfer of SFPs involved in Ovulin and SP networks (long-term exposure / experiment 2 under high-SCR). log_e_ fold-change estimates of protein abundance induced by low (20°C) and hot (28°C) temperatures relative to the 24°C baseline. A) Posterior estimates for SFP production. B) Posterior estimates for SFP transfer. Blue intervals represent the 20°C vs 24°C contrast, while red intervals represent the 28°C vs 24°C contrast. Central point on each segment denotes the median posterior estimate for a specific protein. Thick and thin horizontal lines represent the 50% and 95% Credible Intervals (CrI), respectively. Vertical dashed line at zero represents no change in abundance compared to the 24°C baseline. Estimates where the 95% CrI strictly excludes zero indicate a highly probable (>95%) robust difference induced by the specific temperature treatment.

## Discussion

In this study, we show that temperature variation, within the optimal reproductive range for a wild *D. melanogaster* population, affects both the production and transfer of SFPs, with effects shaped by treatment duration. First, our results show that SFP production and transfer are very plastic in response to temperature. In particular, long-term (13d) exposure to both low (20°C) and high (28°C) temperatures resulted in lower production and transfer of SFPs. Second, we show that these effects carry over to proteins with known important functions in female post-mating responses, whereby long-term exposure to high and low temperatures reduced the production and transfer of SFPs involved in the SP and ovulin networks. Overall, these findings highlight the role of natural temperature fluctuations in shaping ejaculate composition, with potential implications for the dynamics of sexual selection, sexual conflict, and temperature effects on reproduction at large.

### SFP production and transfer are thermally plastic

In line with our predictions, we found a pronounced plastic response in SFP production and transfer after long-term exposure to both high and low temperatures, even though both temperatures fall well within the optimal reproductive range for this population in the wild (<u>Londoño-Nieto et al.,</u> <u>2023)</u>. Males kept at 20°C or 28°C produced and transferred lower abundances of SFPs than at 24°C across SCR levels. In the long-term exposure experiment, sexually mature males were exposed to different stable temperatures for 13-days following partial depletion (males can mate consecutively with up to 10 females; <u>Sanghvi et al., 2025)</u> of accessory-glands via repeated matings (Sirot et al., 2009). Thus, our design ensures that males were forced to synthesize new SFPs under our temperature treatments, fulfilling our aim of capturing effects on the production and transfer of new SFPs that would be typical of seasonal effects. Wild *D. melanogaster* flies are likely to live long enough to experience seasonal thermal fluctuations, given that they can survive up to 90 days in controlled conditions (Brischigliaro et al., 2023). Our results thus support the idea that seasonal changes in temperature may impose physiological constraints on the synthesis and/or transfer of SFPs, and fit well with the finding that male-induced harm to females is significantly reduced at 20°C and 28°C (vs. 24°C) in this same population of flies (Londoño-Nieto et al., 2023).

In contrast, our results following short-term exposure to high and low temperatures were less clear and somewhat paradoxical. We again found clear temperature effects on both ejaculate production and transfer, but thermal plasticity was strongly modulated by sociosexual context in a complex way. Under no-SCR, we found higher production and transfer at both low and high temperatures, but these differences disappeared under high-SCR. Our short-term experiment was designed to capture effects on a fine-grained ecological scale (i.e. short intragenerational effects), given that sexually mature males were exposed to different stable temperatures for 48h and flies likely live considerably longer lifespans in the wild. Compared to our long-term treatment, we may thus expect temperature effects on the production of SFPs to be relatively limited in this treatment.

Broadly, our findings align with previous work showing that high and low temperatures influence several stages of protein synthesis, including transcription, translation, folding, and post-translational modification, which collectively affect protein functionality (Feller & Gerday, 2003; Zheng et al., 2024). In particular, high temperatures are known to be especially detrimental to protein and sperm phenotype and function across animal taxa (Dougherty et al., 2024; Reinhardt et al., 2015; Sales et al., 2018). For instance, heat stress commonly reduces sperm production, motility, viability, and longevity (Wang & Gunderson, 2022). It also increases molecular entropy, which can interfere with correct protein folding and decreases the proportion of functional proteins (Berger et al., 2021). Recent work has extended these findings to seminal fluid components, suggesting that SFPs may also be thermally sensitive (Canal Domenech & Fricke, 2022; Martinet et al., 2023). Our findings build on this work by showing that non-stressful thermal variation within the optimal reproductive range can alter SFP production and transfer. The finding that even subtle and relatively fine-grained temperature variation can have clear effects on SFP production and transfer opens up a new avenue to understand how ecology affects sexual selection, and may also shed light on expected effects of climate change on populations. There is growing recognition that fertility limits, rather than lethal temperature limits, may be more accurate predictors of extinction vulnerability (Dougherty et al., 2024). For instance, some *Drosophila* species can become sterile at temperatures well below their lethal limit (Parratt et al., 2021; Van Heerwaarden & Sgrò, 2021). Our results suggest that, even below these fertility thresholds, natural temperature fluctuations may still influence sexual selection dynamics, potentially increasing vulnerability to climate change in ways not yet fully appreciated (García-Roa et al., 2020).

### Long-term exposure to low and high temperatures reduces the production and transfer of SFP that mediate key post-mating processes

Beyond the general effects of temperature on SFPs plasticity, we predicted that prolonged exposure to 28°C would particularly impact functionally important SFPs involved in mediating female receptivity (Londoño-Nieto et al., 2023). Specifically, Londoño-Nieto et al. (2023) showed that ejaculates from males exposed during 13 days to 28°C induced a markedly weaker reduction in female receptivity (i.e. shorter remating latency) compared to males kept at 20°C and, especially, at 24 °C. On a multivariate level, our results fit well with this finding, as, after 13 days of exposure, we observed consistently lower production and transfer of SP -and ovulin-associated proteins at both 28°C and 20°C compared to 24°C (Fig. 2). In addition, targeted Bayesian multivariate analysis clearly showed a systematic suppression of production and transfer of SPFs in the ovulin and SP networks at 20°C and 28°C vs. 24°C which was particularly marked in males under high-SCR, strongly suggesting that males primed for sperm competition are more sensitive to long-term changes in temperature. Distinguishing whether thermal variation alters the quantity of SFPs produced and transferred versus preserving quantity but compromising function is critical to understand how temperature modulates SFPs dynamics. Both mechanisms would impact male ability to influence female post-mating responses, but they imply different underlying processes and evolutionary consequences. Reduced SFP production and transfer would reflect constraints on male allocation to ejaculates, whereas changes in protein function indicate its effectiveness is thermally constrained, independent of male investment. Our results suggest that, in addition to plasticity in SFP production and transfer, temperature also may affect the functional efficacy of these proteins once transferred.

Ejaculate composition is shaped by trade-offs between ejaculate components that mediate post-mating sexual interactions and sperm competition (Perry et al., 2013). Our results thus beg the question of whether temperature may affect maleś ability to strategically adjust the production and transfer of SFPs in response to perceived SCR, where higher investment is typically observed under high-SCR vs. no-SCR conditions (Hopkins et al., 2019; Wigby et al., 2009). Although our design was not optimized to test this interaction directly, we performed visual comparisons across temperature treatments to explore potential trends in production and transfer of SFPs by males kept under high-SCR vs. no -SCR. We found a trend in that high temperatures seem to disrupt the expected SCR-related patterns of SFP production and transfer in both experiments, especially in SP and ovulin network proteins (Fig. S9). At 28°C, after short-term exposure, males under high-SCR produced relatively less of the proteins belonging to the SP and ovulin networks (including ovulin but excluding SP itself) compared to males under no-SCR, than they did at 24°C, and transferred less *lectin-46Ca, lectin-46Cb, CG9997,* and *SP* than those under no-SCR (Fig. S9A). After prolonged thermal exposure at 28°C, males under high-SCR produced less *Semp1, lectin-46Cb, lectin-46Ca, intr, Sems, antr, CG17575* and *SP*, and transferred less *Sems, lectin-46Ca*, *antr*, *Semp1, ovulin* and *SP* than males under no-SCR, than they did at 24°C (Fig. S9B). This trend must be taken with extreme caution as it is exploratory and preliminary, but it suggests that future studies should address the idea that high temperatures may limit males’ ability to strategically adjust their ejaculate composition in response to SCR.

### The ecological relevance of thermal effects on SFPs

Taken together, our results underscore the broader implications of thermal sensitivity in ejaculate composition. Due to the importance of SFPs, their plasticity (or lack thereof) under thermal variation could mediate individual fitness and shape population responses to environmental change. Our results from experiment 1 suggest that short term fine-scale temperature fluctuations (e.g. circadian) are unlikely to have major effects on females and populations, especially under high-SCR. However, our results from experiment 2 capture the cumulative impact of thermal variation on ejaculate production and transfer over a timeframe that spans both intra- and inter-generational seasonal effects. Such thermal fluctuations are extremely common in the field (particularly in temperate climates), and thus are of clear ecological relevance.

At the intragenerational scale, sustained temperature effects on continuous SFP production and transfer are bound to alter male reproductive dynamics and overall reproductive success across an individual’s lifetime, potentially making plasticity itself a target of selection. Supporting this idea, there is evidence showing that seminal fluid composition can vary among genotypes in response to environmental conditions, with direct consequences for reproductive success (Patlar & Ramm, 2020). In addition, *D. melanogaster* from the same population used in this study show strong thermal GxE interactions in male reproductive success (Londoño-Nieto et al., 2025). Interestingly, these GxE effects appear to be sexually dimorphic (Panyam et al., unpublished results), indicating that males and females may experience, and potentially adapt to, thermal variation in different ways. Moreover, a recent experimental evolution experiment shows the thermal environment can drive quick evolutionary changes in SFPs composition (Londoño-Nieto et al., 2025).

At the intergenerational scale, prolonged periods of heat or cold may shift the targets of post-copulatory sexual selection across seasons, potentially driving seasonal adaptive tracking and/or balancing selection (Bergland et al., 2014; Hoffmann et al., 2005; Machado et al., 2021; Rudman et al., 2022). For example, if higher temperatures consistently reduce the abundance or effectiveness of key proteins like SP, this could weaken male ability to manipulate female post-mating behaviour, potentially shifting the relative importance of pre- vs. post-copulatory sexual selection towards the former. As climate change increases the frequency and magnitude of temperature fluctuations, such effects may be increasingly important for temperate populations of ectothermic species.

## Conclusions

Our results show that natural temperature fluctuations can have a strong effect on SFP production and transfer, and thus on post-copulatory sexual selection dynamics. Notably, long-term exposure to low and high temperatures consistently reduced the production and transfer of SFPs, including a systematic down-regulation of key proteins involved in female post-mating responses (i.e. ovulin and SP networks). These findings highlight how seasonal environmental variation can shape ejaculate traits under realistic ecological conditions, and provide a mechanistic link for effects shown in recent studies associating habitat complexity, reproductive plasticity, and eco-evolutionary processes shaping sexual selection (Berger & Liljestrand-Rönn, 2024; Gomez-Llano et al., 2018; Londoño-Nieto et al., 2023, 2025; Malek & Long, 2019; Yun et al., 2017). Importantly, these results may have important implications for understanding species vulnerability to climate change. We suggest a focus of future studies should be to examine how temperature influences the male’s ability to tailor SFP production and transfer in response to SCR, as well as the plasticity in female reproductive traits (i.e. their responses to male ejaculates), and how these female traits co-evolve with male ejaculates in response to temperature variation. Finally, these findings may also have applied relevance, as the thermal sensitivity of SFPs could improve strategies in pest control and assisted reproduction.

## Author Contributions

Claudia Londoño-Nieto: Conceptualization, data collection and curation, formal analysis, investigation, methodology, writing; Roberto García-Roa: Conceptualization, supervision, investigation, methodology, data collection – supporting, writing-review; Stuart Wigby: Conceptualization, writing-review; Irem Sepil: Conceptualization, writing-review; Pau Carazo: Conceptualization, resources, supervision, funding acquisition, validation, investigation, methodology, data collection - supporting, writing - review and editing.

