## Supplementary Information for "Chilling out or heating up? Thermal plasticity of seminal fluid proteins in *Drosophila melanogaster*"

**Materials and Methods**

Proteomics assays

*Proteomics sample preparation*

Protein extraction and preparation of the SWATH experiment (library and samples) were carried out in the proteomics laboratory of the University of Valencia, Spain, according with the procedure indicated below.

Total protein extracts were prepared by centrifugation of each sample at 13000 rpm 15 min. Supernatants were discarded and pellets suspended in 50 µL of Laemmli buffer 1.5 X. Vortex 5 min and sonicated 5 min. Total protein concentration was calculated using Machery Nagel kit. To prepare library and each sample for SWATH experiment appropriate volume of sample (7.5 µg/sample to SWATH and 25 µg of mixed samples to perform library) was denatured at 95°C during 5 min.

*Spectral Library Building*

Aliquots with an equivalent amount of a selection of samples were mixed to make a pool for building the spectral library (25 µg). The library electrophoresis was performed using a 12% precast gel (Bio-Rad) at 200V for 30 min. Gels were fixed with 40% ethanol/10% acetic acid for one hour and stained with colloidal Coomassie (Bio-Rad) for 15 min. Gels were destained with $H_2O$ milliQ and cutted into six pieces for protein digestion.

In gel protein digestion

The career corresponding to the library was cutted into 6 pieces and then was digested with sequencing grade trypsin (Promega) as described by Shevchenko et al., 1996. 500 ng of trypsin were used for each sample, and digestion was set to 37ºC on. Trypsin digestion was stopped with 10% TFA, the SN was removed, and the library gel slides were dehydrated with pure ACN(Shevchenko et al., 1996). The new peptide solutions were combined with the corresponding SN. The peptide mixtures were dried in a speed vacuum and re suspended in 2% ACN; 0.1% TFA (15 µL) before LC-MS/MS (Liquid chromatography and tandem mass spectrometry/mass spectrometry) analysis.

*LC-MS/MS analysis*

Peptides were analysed using an Ekspert nanoLC 425 nanoflow system (Eksigent Technologies, ABSCIEX) coupled to a mass spectrometer nanoESI qQTOF MS (6600 plus TripleTOF, ABSCIEX). 5 µl of peptide mixture sample was loaded onto a trap column (3µ C18-CL, 350 µm x 0.5 mm; Eksigent) and desalted with 0.1% TFA at 5 µl/min during 5 min. Peptides were then loaded onto an analytical column (3 µ C18-CL 120 Ă, 0.075 x 150 mm; Eksigent) equilibrated in 5% acetonitrile 0.1% FA (formic acid). Elution was carried out with a linear gradient of 7 to 40% B in A for 120 min. (A: 0.1% FA; B: ACN, 0.1% FA) at a flow rate of 300 nL/min. Samples were ionized in a Source Type: Optiflow < 1 µl Nano applying 3.0 kV to the spray emitter at 200 ºC. Analysis was carried out in a data-dependent mode. Survey MS1 scans were acquired from 350–1400 m/z for 250 ms. The quadrupole resolution was set to 'LOW' for MS2 experiments, which were acquired 100–1500 m/z for 25 ms in 'high sensitivity' mode. Following switch criteria were used: charge: 2+ to 4+; minimum intensity; 250 counts per second (cps). Up to 100 ions were selected for fragmentation after each survey scan. Dynamic exclusion was set to 15 s. The rolling collision energies equations were set for all ions as for 2+ ions according to the following equations: |CE|=(slope)x(m/z)+(intercept). The system sensitivity was controlled by analysing 500 ng of K562 trypsin digestion (Sciex). The system sensitivity was controlled with 2 fmol PepCalMix (LC Packings).

*Protein Identification*

ProteinPilot default parameters were used to generate peak list directly from 6600 TripleTof wiff files. The Paragon algorithm (Shilov et al., 2007) of ProteinPilot v 5.0 search engine (ABSciex) was used to search the Uniprot_insecta and Uniprot_Drosophila database with the following parameters: Trypsin specificity, IAM cys-alkylation and the search effort set to through and FDR correction.

The protein grouping was done by Pro group algorithm: A protein group in a Pro Group Report is a set of proteins that share some physical evidence. Unlike sequence alignment analyses where full-length theoretical sequences are compared, the formation of protein groups in Pro Group is guided entirely by observed peptides only. Since the observed peptides are determined from experimentally acquired spectra, the grouping can be considered to be guided by usage of spectra. Then, unobserved regions of protein sequence play no role in explaining the data.

*SWATH analysis of individual samples*

For individual SWATH analysis 7.5 µg of total protein extract was loaded in a 1D_SDS_PAGE gel to clean and concentrate samples. Gel fraction was cut and the sample was digested with sequencing grade trypsin (Promega) as described elsewhere (Shevchenko et al., 1996). 500 ng of trypsin in 100 µl of ABC solution was used. The digestion was stopped with TFA (1% final concentration), a double extraction with ACN was done and all the peptide solutions and dried in a rotatory evaporator. Sample was re suspended with 15 µL of 2% ACN; 0.1% TFA.

*SWATH LC-MS/MS Analysis*

5 µl of each sample were loaded onto a trap column (3µ C18-CL 120 Ă, 350 µm x 0.5mm; Eksigent) and desalted with 0.1% TFA at 5 µl/min during 5 min. Peptides were loaded onto an analytical column (3µ C18-CL 120 Ă, 0.075 x 150 mm; Eksigent) equilibrated in 5% acetonitrile 0.1% FA (formic acid). Peptide elution was carried out with a linear gradient of 7 to 40% B in 120 min (A: 0.1% FA; B: ACN, 0.1% FA) for at a flow rate of 300 nl/min. Peptides were analysed in a mass spectrometer nanoESI qQTOF (6600plus TripleTOF, ABSCIEX).

Sample was ionized in a Source Type: Optiflow < 1 µl Nano applying 3.0 kV to the spray emitter at 200ºC. The tripleTOF was operated in swath mode, in which 0.050-s TOF MS scan from 350–1250 m/z was performed, followed by 0.080-s product ion scans from 350–1250 m/z. 100 variable windows from 400 to 1250 m/z were acquired throughout the experiment. The total cycle time was 2.79 secs. The individual SWATH injections were randomized.

*Protein quantification*

The wiff files obtained from SWATH experiment were analysed by Peak View 2.2. The processing settings used for the peptide selection were: 20 peptides per protein, 6 transitions per peptide, 95% peptide confidence threshold, 1.0% false discovery rate threshold, peptides modified excluded, 5 min XIC extraction window and 25 ppm XIC width.

Retention times of the detected peptides were alienate using major proteins to calibrate retention times. With the extraction parameters of the areas used, proteins (FDR <1%) were quantified in the 72 samples.

*Data analysis*

The optimal value of $\alpha$ and $\lambda$, estimated using cross-validation (Zou & Hastie, 2005), for our analyses are in the Table S1.

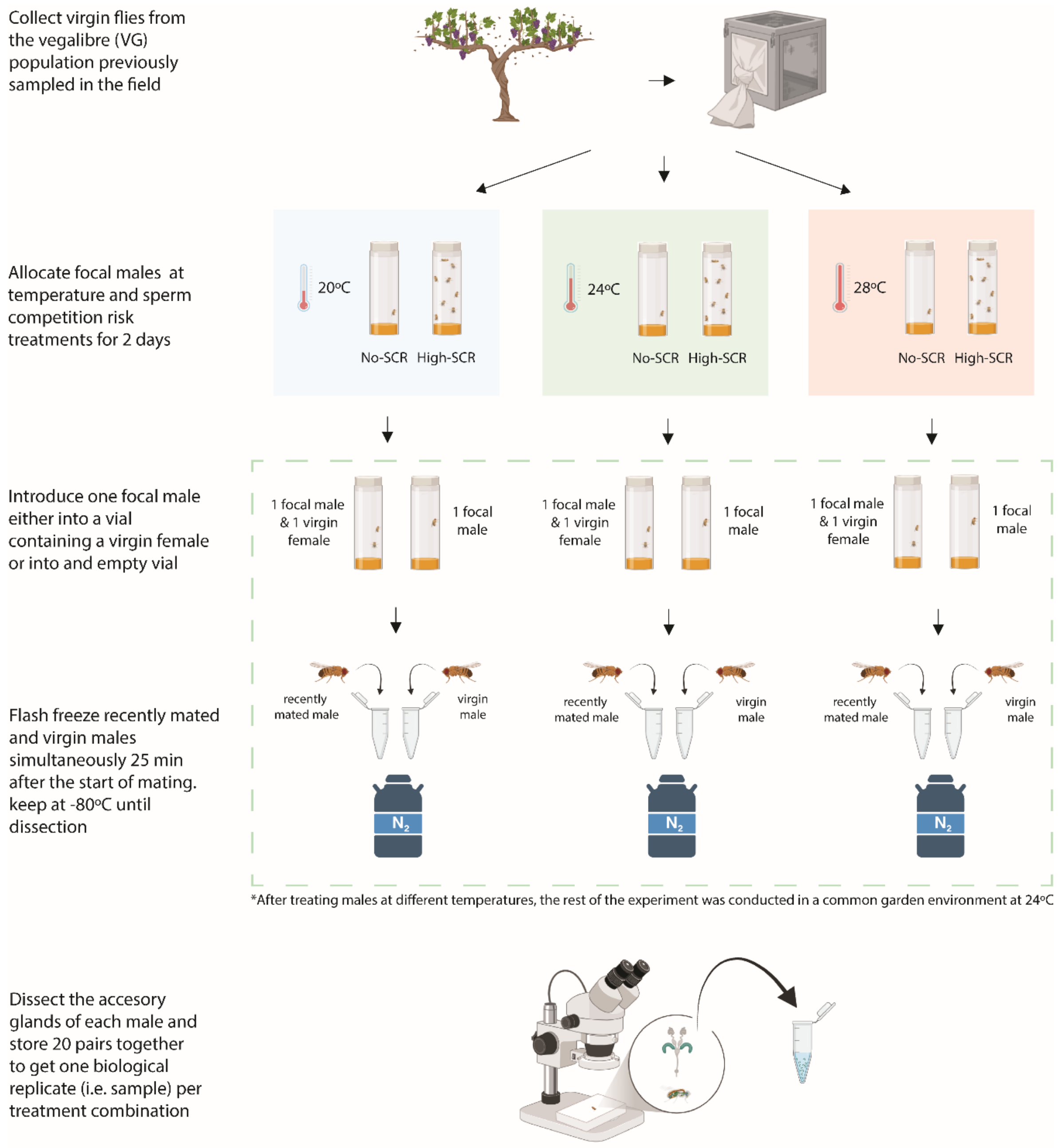


**Figure S1.** Proteomics assay design, experiment 1 (48-hour treatment duration)

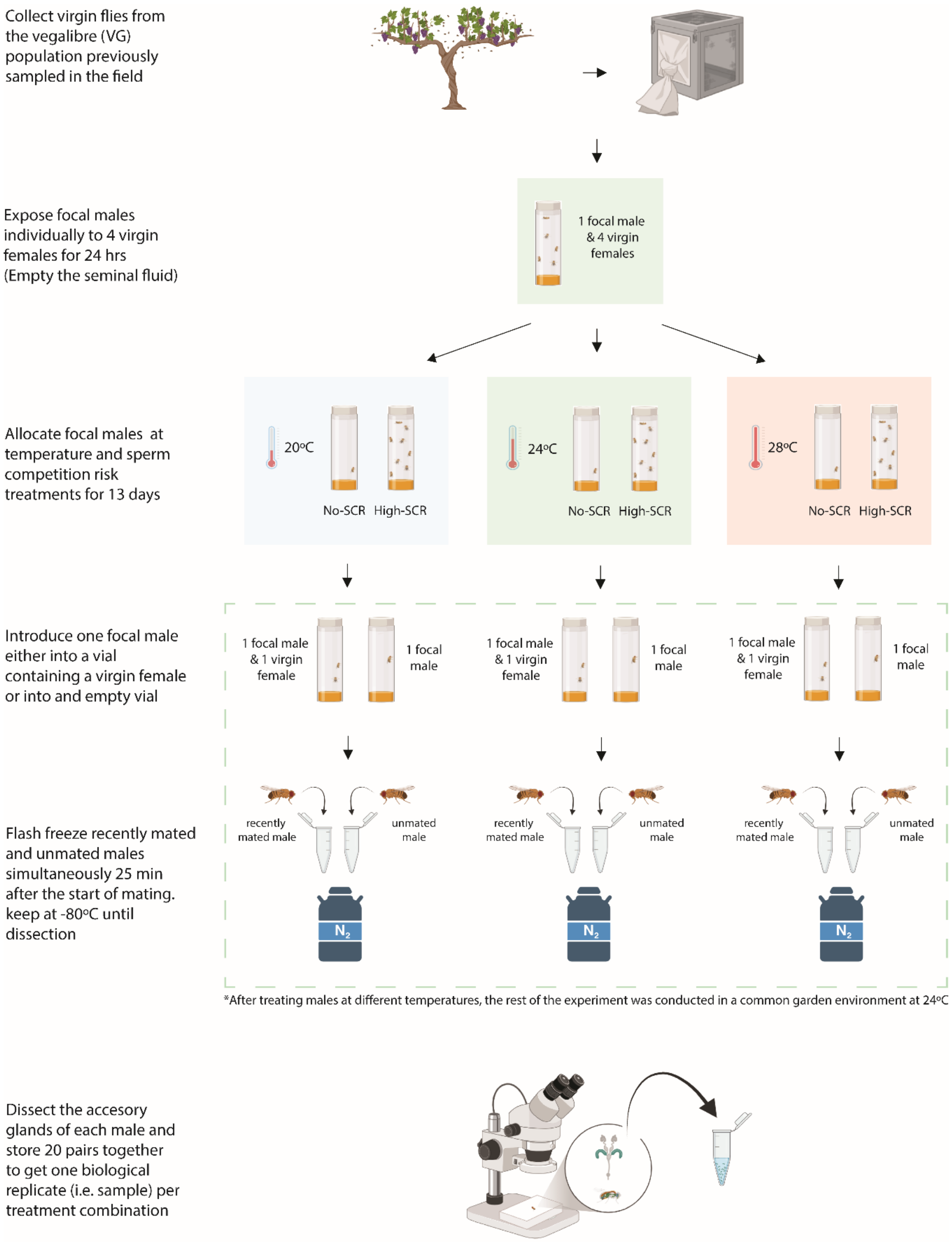


**Figure S2.** Proteomics assay design, experiment 2 (13-days treatment duration)

### Results

**Table S1.** Parameters used for Elastic net analysis. Shaded datasets correspond to those used to analyse **SFP production**, while unshaded datasets correspond to those used to analyse **SFP transfer** for each experiment.

| Experiment | Data set | α | λ |
|---|---|---|---|
| 1 (48hours) | Virgin males (no SCR) | 0,1 | 1,628 |
| | Virgin males (high SCR) | 1 | 0,280 |
| | Virgin/Newly mated ratio (no SCR) | 0,1 | 0,123 |
| | Virgin/Newly mated ratio (high SCR) | 1 | 0,082 |
| 2 (13 days) | Unmated males (no SCR) | 0,4 | 0,095 |
| | Unmated males (high SCR) | 0,1 | 1,297 |
| | Unmated/Newly mated ratio (no SCR) | 0,5 | 0,015 |
| | Unmated/Newly mated ratio (high SCR) | 0,1 | 2,499 |

**Table S2**. Coefficients of 74 proteins selected by elastic net regression as predictors with the strongest effects on proteome quantification for **SFP production under no-SCR in Experiment 1** (short-term exposure). Non-zero coefficients indicate proteins retained by the model and considered to contribute most strongly to variation in protein abundance across temperature treatment.

| Protein | Coefficients | | |
|---|---|---|---|
| | 20ºC | 24ºC | 28ºC |
| X2J8Y6 | -0,01336 | 0,010356 | 0,003002 |
| Q9VAY2 | 0,003181 | -0,00784 | 0,004655 |
| X2JGP4 | 3,15E-05 | -8,9E-05 | 5,76E-05 |
| Q9U1I7 | 0,006207 | -0,00136 | -0,00485 |
| O46067 | 0,011656 | -0,00905 | -0,0026 |
| Q7K110 | -0,00092 | -0,0019 | 0,00282 |
| C9QP21 | 0,027092 | -0,01192 | -0,01517 |
| Q9W0F7 | 0,000632 | -0,00929 | 0,00866 |
| Q6GUS0 | -0,0176 | -0,03215 | 0,049746 |
| Q9VLB4 | 0,00233 | -0,00533 | 0,002996 |
| O46199 | 0,042304 | -0,01004 | -0,03226 |
| Q9VG48 | 0,010277 | -0,01674 | 0,006464 |
| Q9VIL5 | 0,018541 | 0,030317 | -0,04886 |
| Q2PDZ4 | 0,008025 | -0,01176 | 0,00373 |
| Q8T429 | 0,014226 | -0,02183 | 0,007603 |
| A0A0B4LGZ1 | 0,013024 | -0,01839 | 0,005368 |
| Q7YTY6 | 0,009722 | -0,00934 | -0,00039 |
| Q7KMM4 | 0,010177 | -0,01498 | 0,004799 |
| Q9VWT3 | 0,022122 | -0,0215 | -0,00062 |
| Q95S79 | 0,011431 | -0,00444 | -0,00699 |
| Q9W3C4 | -0,00575 | -0,00493 | 0,010679 |
| A1Z957 | 0,000803 | -0,00244 | 0,001642 |
| Q9W227 | -0,01313 | -0,0036 | 0,016737 |
| Q7JY62 | 0,012872 | -0,01227 | -0,0006 |
| Q9VHH1 | 0,000845 | -0,00174 | 0,000891 |
| Q9VLV3 | 0,01524 | -0,0058 | -0,00944 |
| A1Z959 | 0,002869 | -0,00233 | -0,00054 |
| Q9VP92 | -0,01963 | 0,027578 | -0,00795 |
| Q9VVQ6 | 0,006368 | -0,02317 | 0,016802 |
| Q9W478 | 0,006696 | -0,01126 | 0,004569 |
| Q9V3Q4 | 0,04704 | -0,03028 | -0,01676 |
| Q9VP95 | 0,023758 | -0,03219 | 0,008437 |
| Q9VGA5 | 0,004537 | -0,00436 | -0,00018 |
| Q7KE32 | -0,03775 | -0,02519 | 0,062946 |
| Q4V6H2 | 0,022732 | -0,00945 | -0,01328 |
| Q9VX69 | 0,012525 | -0,00813 | -0,00439 |
| Q9VY09 | 0,01402 | -0,02831 | 0,014292 |
| Q9VBG5 | 0,011918 | -0,01259 | 0,000671 |
| Q0KHY7 | 0,004187 | -0,02315 | 0,018963 |
| Q9VGQ4 | 0,036765 | -0,01875 | -0,01801 |
| A8DY52 | -0,00808 | -0,02198 | 0,030053 |
| Q9VAX6 | 0,015835 | -0,0205 | 0,004665 |
| Q9VPH9 | 0,003583 | -0,00166 | -0,00192 |
| Q9VPS2 | 0,034909 | -0,05182 | 0,016911 |
| A0A0B4K830 | 0,01254 | -0,00998 | -0,00256 |
| Q9V3J7 | -0,00632 | -0,02371 | 0,030029 |
| E2QCL8 | 0,007649 | -0,02653 | 0,01888 |
| X2JEB6 | -0,02873 | 0,006901 | 0,021828 |
| Q7K088 | -0,00408 | -0,02512 | 0,029204 |
| Q9VI09 | -0,03309 | -0,02036 | 0,053459 |
| Q9VHJ0 | 0,019358 | -0,01974 | 0,000385 |
| Q8MSK0 | -0,02496 | -0,02432 | 0,049276 |
| Q9VDI5 | 0,007491 | -0,00744 | -4,7E-05 |
| Q6IGA5 | 0,020138 | -0,01757 | -0,00257 |
| C0PV13 | -0,02795 | -0,00199 | 0,029934 |
| D1Z365 | 0,004269 | -0,03734 | 0,03307 |
| Q9VXU9 | 0,002526 | -0,00431 | 0,001788 |
| Q9XZ54 | 0,019874 | -0,041 | 0,021126 |
| Q95SR0 | 0,002005 | -0,00143 | -0,00058 |
| Q9VWV6 | 0,019861 | -0,00961 | -0,01026 |
| Q6GUX9 | 0,006886 | -0,00871 | 0,001828 |
| A8JRF1 | 9,13E-05 | -0,00012 | 2,57E-05 |
| Q9VJN9 | -0,00165 | -0,00052 | 0,002173 |
| Q9VR13 | -0,00339 | 0,001052 | 0,002335 |
| A8DY51 | 0,008797 | 0,005016 | -0,01381 |
| Q9VCM5 | 0,000816 | -0,00089 | 7,31E-05 |
| Q9VFN7 | 0,029689 | -0,0417 | 0,012014 |
| Q8MS62 | 0,01396 | -0,02261 | 0,008653 |
| Q9VZ58 | -0,02421 | -0,01998 | 0,044191 |
| B4ZJA4 | -0,01824 | -0,00367 | 0,021908 |
| Q9V3J3 | 0,029657 | 0,033823 | -0,06348 |
| Q6GUV0 | 0,00272 | -0,00837 | 0,005646 |
| O61457 | 0,004405 | -0,00929 | 0,004886 |
| Q4V3K7 | 0,002726 | 0,007355 | -0,01008 |

**Table S3**. Coefficients of 3 proteins selected by elastic net regression as predictors with the strongest effects on proteome quantification for **SFP production under High-SCR in Experiment 1** (short-term exposure). Non-zero coefficients indicate proteins retained by the model and considered to contribute most strongly to variation in protein abundance across temperature treatment.

| Protein | Coefficients | | |
|---|---|---|---|
| | 20ºC | 24ºC | 28ºC |
| E1JHG2 | -1,05658047 | 0,60526451 | 0,45131596 |
| Q9V3J3 | 0,32725288 | -0,23315475 | -0,09409814 |
| O61457 | -0,06964846 | -0,32926141 | 0,39890987 |

**Table S4**. Coefficients of 79 proteins selected by elastic net regression as predictors with the strongest effects on proteome quantification for **SFP transfer under No-SCR in Experiment 1** (short-term exposure). Non-zero coefficients indicate proteins retained by the model and considered to contribute most strongly to variation in protein abundance across temperature treatment.

| Protein | Coefficients | | |
|---|---|---|---|
| | 20ºC | 24ºC | 28ºC |
| Q9VKT1 | 0,004304 | 0,040110 | -0,044414 |
| Q9U1I7 | 0,009539 | 0,011308 | -0,020847 |
| Q9VD61 | -0,001322 | -0,000898 | 0,002220 |
| Q7K110 | -0,004832 | -0,005515 | 0,010347 |
| Q9VAT7 | 0,017749 | -0,006010 | -0,011739 |
| C9QP21 | 0,010895 | -0,000368 | -0,010527 |
| P10333 | 0,011169 | -0,015051 | 0,003882 |
| Q3HKQ3 | 0,003671 | -0,002043 | -0,001628 |
| Q8MLV0 | -0,015852 | 0,023628 | -0,007775 |
| Q9VLZ8 | 0,019856 | -0,016799 | -0,003057 |
| Q9VG48 | 0,031762 | -0,031938 | 0,000176 |
| Q9VG47 | 0,003067 | -0,002372 | -0,000695 |
| Q9VIL5 | 0,008264 | 0,027104 | -0,035369 |
| Q9VAT8 | 0,013847 | -0,011699 | -0,002148 |
| Q2PDZ4 | 0,016777 | -0,021590 | 0,004813 |
| Q8T429 | 0,002105 | -0,002126 | 0,000021 |
| A0A0B4LGZ1 | 0,006681 | -0,008903 | 0,002222 |
| Q7YTY6 | 0,006062 | -0,003516 | -0,002546 |
| Q95S79 | 0,015977 | -0,010579 | -0,005398 |
| Q8IPX4 | 0,012023 | -0,010665 | -0,001358 |
| Q6GV06 | 0,004023 | -0,004712 | 0,000689 |
| Q9W129 | 0,024564 | -0,019287 | -0,005276 |
| Q0KI07 | 0,009544 | -0,004470 | -0,005074 |
| Q7JY62 | 0,014607 | -0,011446 | -0,003161 |
| Q6IH48 | 0,003088 | -0,001890 | -0,001198 |
| Q9VVQ5 | 0,002775 | -0,007777 | 0,005002 |
| Q8MVX6 | 0,027079 | 0,009780 | -0,036858 |
| Q9VP92 | -0,009597 | 0,023181 | -0,013583 |
| Q8T971 | 0,017042 | 0,026396 | -0,043439 |
| Q9VVQ6 | 0,005763 | -0,017669 | 0,011906 |
| Q9VEL5 | 0,003525 | -0,001897 | -0,001628 |
| Q9V3Q4 | 0,023265 | -0,026418 | 0,003153 |
| Q9VP95 | 0,001522 | -0,000748 | -0,000773 |
| Q9VGA5 | 0,014113 | -0,014059 | -0,000053 |
| Q4V6H2 | 0,000045 | -0,000010 | -0,000035 |
| Q6IGS3 | 0,014167 | -0,006081 | -0,008086 |
| Q9VX69 | 0,007741 | 0,006603 | -0,014344 |
| Q9VY09 | -0,007354 | -0,035353 | 0,042707 |
| E1JJ84 | 0,000466 | 0,000550 | -0,001017 |
| Q9VBG5 | 0,005574 | -0,003880 | -0,001693 |
| Q4V566 | 0,006682 | -0,007245 | 0,000563 |
| A8DY52 | -0,006953 | -0,012859 | 0,019811 |
| Q9VAX6 | 0,010087 | -0,012349 | 0,002262 |
| Q9VPH9 | 0,008281 | -0,004789 | -0,003492 |
| Q9VPS2 | 0,016194 | -0,013031 | -0,003163 |
| Q6IHT7 | 0,006716 | 0,006567 | -0,013283 |
| Q9VKS9 | 0,001417 | -0,001167 | -0,000250 |
| Q9VS71 | -0,007062 | -0,009974 | 0,017036 |
| E2QCL8 | 0,012752 | -0,022000 | 0,009248 |
| X2JEB6 | -0,022358 | 0,008692 | 0,013666 |
| M9NEQ6 | 0,034202 | -0,024891 | -0,009310 |
| Q9VAT6 | 0,026271 | 0,005476 | -0,031748 |
| E1JHS6 | 0,000947 | 0,000203 | -0,001151 |
| E1JHF8 | 0,004156 | -0,000120 | -0,004036 |
| F0JAR5 | 0,003131 | 0,021288 | -0,024419 |
| Q3HKQ0 | 0,001168 | -0,000372 | -0,000796 |
| Q6GV01 | 0,001813 | -0,007030 | 0,005216 |
| Q9VI09 | -0,031887 | -0,022016 | 0,053903 |
| Q9VHJ0 | 0,024517 | -0,019272 | -0,005244 |
| Q9VDI5 | 0,008774 | -0,007596 | -0,001179 |
| Q6IGA5 | 0,006982 | -0,000508 | -0,006474 |
| D1Z365 | 0,011313 | -0,020981 | 0,009668 |
| Q9XZ54 | 0,001621 | -0,009977 | 0,008356 |
| Q95SR0 | 0,002219 | -0,002385 | 0,000166 |
| Q9VWV6 | 0,006021 | 0,010075 | -0,016095 |
| Q6GUX9 | 0,010634 | -0,007811 | -0,002823 |
| Q8T4B0 | -0,090953 | 0,126015 | -0,035061 |
| A8JRF1 | 0,035764 | 0,048161 | -0,083926 |
| Q9VR13 | -0,042454 | 0,023172 | 0,019282 |
| Q9VCM5 | 0,004386 | -0,006249 | 0,001863 |
| Q9VLQ7 | 0,005253 | -0,002925 | -0,002328 |
| Q3HKQ1 | 0,000276 | -0,003320 | 0,003044 |
| Q9VZ58 | -0,009412 | -0,000981 | 0,010393 |
| Q9V3J3 | 0,003751 | 0,000582 | -0,004333 |
| Q6GUV0 | -0,001991 | 0,007734 | -0,005743 |
| Q0KI39 | 0,012593 | -0,011933 | -0,000660 |
| O61457 | 0,007097 | -0,009274 | 0,002177 |
| Q4V3K7 | -0,006870 | -0,010158 | 0,017028 |
| Q9W2S7 | 0,037486 | 0,007738 | -0,045224 |

**Table S5**. Coefficients of 6 proteins selected by elastic net regression as predictors with the strongest effects on proteome quantification for **SFP transfer under High-SCR in Experiment 1** (short-term exposure). Non-zero coefficients indicate proteins retained by the model and considered to contribute most strongly to variation in protein abundance across temperature treatment.

| Protein | Coefficients | | |
|---|---|---|---|
| | 20ºC | 24ºC | 28ºC |
| Q8MR48 | -0,134266 | -0,073655 | 0,207921 |
| E1JHG2 | -0,530915 | 0,933227 | -0,402312 |
| C0PV13 | -0,803587 | 0,452193 | 0,351393 |
| Q9VWV6 | -0,000355 | 0,000381 | -0,000026 |
| Q9VCM5 | -0,151188 | 0,428040 | -0,276852 |
| O61457 | -0,106878 | -0,188214 | 0,295093 |

**Table S6**. Coefficients of 39 proteins selected by elastic net regression as predictors with the strongest effects on proteome quantification for **SFP production under No-SCR in Experiment 2** (long-term exposure). Non-zero coefficients indicate proteins retained by the model and considered to contribute most strongly to variation in protein abundance across temperature treatment.

| Protein | Coefficients | | |
|---|---|---|---|
| | 20ºC | 24ºC | 28ºC |
| C9QP21 | -0,017555 | 0,042322 | -0,024768 |
| Q3HKQ3 | -0,004591 | 0,003170 | 0,001422 |
| Q9VG47 | -0,019207 | 0,023849 | -0,004642 |
| Q9VWB4 | -0,121790 | 0,179515 | -0,057725 |
| Q9W227 | -0,219661 | -0,021398 | 0,241059 |
| Q0KI07 | -0,011413 | 0,008923 | 0,002490 |
| Q7KE32 | -0,145075 | -0,138557 | 0,283632 |
| Q4V6H2 | -0,072121 | 0,165449 | -0,093328 |
| Q9VQA3 | -0,034946 | 0,135592 | -0,100647 |
| E1JJ84 | -0,023347 | 0,071100 | -0,047753 |
| Q9VBG5 | -0,051810 | 0,070620 | -0,018810 |
| E1JHG2 | -0,217335 | -0,183537 | 0,400872 |
| Q6IGA4 | -0,168867 | -0,161793 | 0,330660 |
| Q6IHT7 | -0,065224 | 0,028650 | 0,036574 |
| Q9W2S8 | -0,005388 | -0,013687 | 0,019075 |
| Q9VES3 | 0,057397 | -0,068033 | 0,010637 |
| E2QCL8 | -0,052245 | 0,027443 | 0,024802 |
| M9NEQ6 | -0,049525 | 0,015367 | 0,034158 |
| Q7K088 | -0,022924 | -0,012653 | 0,035577 |
| Q9VAT6 | -0,068394 | 0,062066 | 0,006328 |
| E1JHS6 | -0,000859 | 0,000312 | 0,000547 |
| Q8MLS8 | -0,193753 | 0,083643 | 0,110110 |
| E1JHF8 | -0,064692 | 0,081937 | -0,017245 |
| F0JAR5 | -0,039798 | 0,264168 | -0,224370 |
| Q3HKQ0 | -0,151959 | 0,026309 | 0,125650 |
| Q9VI09 | -0,021395 | 0,067806 | -0,046411 |
| Q9VDI5 | -0,124788 | 0,087519 | 0,037269 |
| C4NAP3 | -0,023511 | -0,057433 | 0,080943 |
| D1Z365 | -0,179291 | 0,079499 | 0,099792 |
| Q9VQE1 | -0,130073 | 0,126833 | 0,003240 |
| X2J4U0 | -0,001219 | 0,000045 | 0,001174 |
| Q6GUX9 | 0,000646 | 0,011651 | -0,012296 |
| Q9VR13 | -0,081247 | 0,255394 | -0,174146 |
| Q9VCM5 | -0,032091 | 0,053038 | -0,020946 |
| Q9VFN7 | 0,028128 | -0,187431 | 0,159303 |
| Q3HKQ1 | -0,005239 | 0,008267 | -0,003028 |
| Q6GUV0 | 0,029506 | 0,013155 | -0,042661 |
| Q0KI39 | -0,152935 | 0,122673 | 0,030262 |
| O61457 | -0,103971 | -0,081459 | 0,185430 |

**Table S7**. Coefficients of 68 proteins selected by elastic net regression as predictors with the strongest effects on proteome quantification for **SFP production under High-SCR in Experiment 2** (long-term exposure). Non-zero coefficients indicate proteins retained by the model and considered to contribute most strongly to variation in protein abundance across temperature treatment

| Protein | Coefficients | | |
|---|---|---|---|
| | 20ºC | 24ºC | 28ºC |
| Q9VKT1 | -0,014211 | 0,008289 | 0,005922 |
| Q9VAY2 | -0,024192 | -0,011908 | 0,036100 |
| Q9VTL4 | -0,047424 | -0,004463 | 0,051887 |
| Q9U1I7 | -0,020521 | 0,015615 | 0,004907 |
| C9QP21 | -0,015023 | 0,028809 | -0,013786 |
| Q86BL9 | 0,000024 | 0,000030 | -0,000054 |
| Q6GUS0 | -0,007364 | 0,002052 | 0,005312 |
| Q8MLV0 | -0,007378 | 0,003186 | 0,004191 |
| O46199 | -0,003409 | 0,001383 | 0,002026 |
| Q24238 | -0,042240 | -0,012870 | 0,055110 |
| Q9VYR1 | -0,009665 | 0,023565 | -0,013900 |
| Q9VIP6 | -0,035967 | 0,029407 | 0,006560 |
| M9PG65 | -0,061797 | 0,047100 | 0,014697 |
| Q9VI96 | -0,000487 | 0,001307 | -0,000820 |
| A0A0B4LGZ1 | -0,005059 | 0,046144 | -0,041084 |
| Q7KMM4 | -0,057400 | 0,031709 | 0,025691 |
| A1Z957 | -0,001586 | 0,002070 | -0,000484 |
| Q7K173 | 0,005954 | 0,000073 | -0,006027 |
| Q9W227 | -0,044208 | 0,000598 | 0,043609 |
| Q9VEL4 | -0,012435 | 0,018283 | -0,005848 |
| Q9VHH1 | -0,010808 | 0,013475 | -0,002666 |
| A0A0B4K7N3 | -0,000983 | 0,009108 | -0,008125 |
| Q9VLV3 | 0,011169 | 0,017168 | -0,028337 |
| Q8T971 | -0,034786 | 0,079452 | -0,044666 |
| Q9W478 | -0,001893 | 0,005846 | -0,003953 |
| Q9VEL5 | -0,000652 | 0,003538 | -0,002886 |
| Q9V3Q4 | 0,004392 | 0,005158 | -0,009550 |
| Q7KE32 | -0,088407 | 0,014982 | 0,073425 |
| Q4V6H2 | -0,004379 | 0,009183 | -0,004804 |
| Q6IGS3 | -0,003714 | 0,012204 | -0,008490 |
| Q9VX69 | -0,012718 | 0,042762 | -0,030044 |
| Q9VY09 | -0,001302 | 0,009885 | -0,008582 |
| E1JJ84 | -0,009530 | 0,016515 | -0,006985 |
| E1JHG2 | -0,128306 | 0,044061 | 0,084245 |
| Q4V566 | -0,005845 | 0,004901 | 0,000944 |
| Q9VGQ4 | -0,019833 | 0,023706 | -0,003873 |
| Q9VAX6 | -0,000386 | 0,000899 | -0,000513 |
| Q7KE33 | -0,016146 | 0,067905 | -0,051759 |
| Q6IHT7 | -0,004654 | 0,001216 | 0,003438 |
| Q9VS71 | 0,001792 | 0,011477 | -0,013269 |
| M9NEQ6 | -0,007501 | 0,036224 | -0,028723 |
| Q7K088 | -0,050050 | 0,034330 | 0,015721 |
| E1JHS6 | -0,014904 | 0,030151 | -0,015246 |
| Q3HKQ0 | -0,077471 | -0,004126 | 0,081598 |
| Q9VHJ0 | -0,005731 | 0,044272 | -0,038541 |
| B4ZJA2 | -0,000162 | 0,000144 | 0,000018 |
| Q6IGA5 | 0,005250 | 0,029184 | -0,034433 |
| C0PV13 | -0,016297 | 0,006431 | 0,009866 |
| Q8IQS7 | -0,006762 | 0,040596 | -0,033834 |
| C4NAP3 | -0,050795 | 0,004254 | 0,046540 |
| Q9VXU9 | -0,006487 | 0,009972 | -0,003485 |
| Q9VQE1 | -0,016877 | 0,029578 | -0,012701 |
| X2J4U0 | -0,029474 | -0,005170 | 0,034645 |
| Q9XZ54 | -0,017456 | 0,015245 | 0,002211 |
| Q9VWV6 | -0,007744 | 0,013667 | -0,005923 |
| Q8T4B0 | -0,035919 | 0,008398 | 0,027522 |
| Q9VJN9 | -0,005406 | 0,022275 | -0,016869 |
| Q9VR13 | -0,015169 | -0,000150 | 0,015319 |
| X2J8T8 | -0,075633 | 0,001003 | 0,074630 |
| Q9VCM5 | -0,003842 | 0,005204 | -0,001362 |
| Q3HKQ1 | -0,002612 | 0,004543 | -0,001931 |
| Q8MS62 | -0,065673 | 0,030564 | 0,035109 |
| Q9VZ58 | -0,005770 | 0,061180 | -0,055410 |
| B4ZJA4 | -0,088071 | 0,020116 | 0,067955 |
| Q9V3J3 | -0,003011 | -0,000296 | 0,003307 |
| Q6GUV0 | -0,009697 | -0,014668 | 0,024365 |
| O61457 | -0,029610 | 0,010714 | 0,018896 |
| Q4V3K7 | -0,000218 | 0,000561 | -0,000343 |

**Table S8**. Coefficients of 49 proteins selected by elastic net regression as predictors with the strongest effects on proteome quantification for **SFP transfer under No-SCR in Experiment 2** (long-term exposure). Non-zero coefficients indicate proteins retained by the model and considered to contribute most strongly to variation in protein abundance across temperature treatment

| Protein | Coefficients | | |
|---|---|---|---|
| | 20ºC | 24ºC | 28ºC |
| Q9VKT1 | 0,097142 | -0,086585 | -0,010557 |
| Q9VAY2 | -0,441108 | -0,687232 | 1,128340 |
| Q9VII7 | -0,087257 | 0,128096 | -0,040840 |
| X2JGP4 | -0,025070 | 0,037555 | -0,012485 |
| Q9VD61 | -0,006729 | 0,006239 | 0,000489 |
| Q9VD62 | -0,127035 | 0,131424 | -0,004389 |
| Q8IMY4 | -0,065589 | 0,025595 | 0,039994 |
| P10333 | -0,019910 | 0,083441 | -0,063531 |
| Q3HKQ3 | -0,011619 | 0,005778 | 0,005841 |
| Q8MLV0 | -0,038115 | 0,043457 | -0,005342 |
| Q9VLB4 | -0,002169 | 0,005666 | -0,003497 |
| Q9VG48 | -0,011912 | 0,048743 | -0,036831 |
| M9PG65 | 0,012581 | 0,071008 | -0,083589 |
| Q2PDZ4 | -0,023888 | 0,027332 | -0,003444 |
| Q9VWB4 | -0,005182 | 0,012145 | -0,006963 |
| Q8T429 | -0,045501 | 0,030922 | 0,014579 |
| Q9W3C4 | -0,097868 | 0,091871 | 0,005997 |
| Q8IPX4 | -0,180983 | 0,122812 | 0,058171 |
| A1Z957 | -0,117527 | 0,055120 | 0,062407 |
| Q9W227 | -0,313213 | 0,062910 | 0,250303 |
| Q7JY62 | -0,128754 | 0,011802 | 0,116953 |
| Q9VEL4 | -0,086173 | 0,082312 | 0,003861 |
| A0A0B4K7N3 | 0,092780 | 0,056412 | -0,149192 |
| Q9VLV3 | 0,048885 | 0,105462 | -0,154347 |
| Q8T971 | 0,243249 | 0,111033 | -0,354282 |
| Q9VVQ6 | -0,032655 | 0,048657 | -0,016002 |
| Q9VGA5 | -0,013626 | -0,002445 | 0,016071 |
| Q7KE32 | -0,200811 | -0,080569 | 0,281380 |
| Q9VQA3 | -0,007769 | 0,095422 | -0,087654 |
| Q9VY09 | 0,099775 | -0,047003 | -0,052772 |
| Q9VBG5 | -0,000333 | 0,000926 | -0,000593 |
| E1JHG2 | -0,395231 | -0,035108 | 0,430340 |
| Q7KE33 | 0,066318 | 0,066111 | -0,132430 |
| Q9VES3 | 0,030271 | -0,014027 | -0,016244 |
| X2JEB6 | 0,010176 | 0,006697 | -0,016873 |
| F0JAR5 | -0,014594 | 0,282886 | -0,268293 |
| Q3HKQ0 | -0,311785 | 0,466845 | -0,155060 |
| Q9VI09 | -0,001168 | 0,007517 | -0,006349 |
| Q9VQE1 | -0,193860 | 0,190649 | 0,003212 |
| Q6GUX9 | -0,014418 | 0,044652 | -0,030234 |
| A8JRF1 | 0,031414 | -0,011071 | -0,020343 |
| Q9VJN9 | -0,039680 | 0,009192 | 0,030488 |
| Q9VCM5 | 0,006668 | 0,010966 | -0,017634 |
| Q9VFN7 | -0,248026 | 0,009121 | 0,238905 |
| Q3HKQ1 | -0,014570 | 0,091357 | -0,076787 |
| Q8MS62 | 0,026099 | 0,061543 | -0,087641 |
| Q9V3J3 | 0,094234 | 0,213024 | -0,307258 |
| Q6GUV0 | 0,049045 | 0,007904 | -0,056949 |
| O61457 | -0,049046 | 0,020382 | 0,028665 |

**Table S9**. Coefficients of 46 proteins selected by elastic net regression as predictors with the strongest effects on proteome quantification for **SFP transfer under High-SCR in Experiment 2** (long-term exposure). Non-zero coefficients indicate proteins retained by the model and considered to contribute most strongly to variation in protein abundance across temperature treatment

| Protein | Coefficients | | |
|---|---|---|---|
| | 20ºC | 24ºC | 28ºC |
| X2JGP4 | -0,006003 | 0,007730 | -0,001726 |
| Q9U1I7 | -0,002693 | 0,002883 | -0,000191 |
| Q9VD62 | -0,000473 | 0,000624 | -0,000151 |
| Q7K110 | -0,026687 | 0,020953 | 0,005734 |
| C9QP21 | -0,002865 | 0,007033 | -0,004168 |
| Q8MLV0 | -0,005262 | 0,001687 | 0,003575 |
| O46199 | -0,000080 | 0,000077 | 0,000003 |
| Q9VG48 | -0,002846 | 0,003477 | -0,000632 |
| Q9VIP6 | -0,005110 | 0,003927 | 0,001183 |
| Q2PDZ4 | -0,002413 | 0,006838 | -0,004424 |
| Q9VI96 | -0,000970 | 0,001392 | -0,000423 |
| A0A0B4LGZ1 | -0,000167 | 0,003367 | -0,003200 |
| Q7KMM4 | -0,029604 | 0,026241 | 0,003363 |
| Q8IPX4 | 0,000536 | 0,008593 | -0,009129 |
| A1Z957 | -0,001111 | 0,002445 | -0,001334 |
| Q7JY62 | -0,000390 | 0,006865 | -0,006475 |
| Q8MR48 | -0,008096 | 0,011016 | -0,002920 |
| Q9VEL4 | -0,013934 | 0,031567 | -0,017633 |
| Q9VHH1 | -0,005993 | 0,010771 | -0,004778 |
| Q9VLV3 | -0,000742 | 0,025176 | -0,024434 |
| Q9VVQ6 | -0,002753 | 0,002677 | 0,000075 |
| Q9W478 | -0,004154 | 0,009127 | -0,004973 |
| Q9VEL5 | -0,000277 | 0,002093 | -0,001816 |
| Q9V3Q4 | -0,000914 | 0,006023 | -0,005110 |
| Q7KE32 | -0,006025 | 0,021445 | -0,015420 |
| E1JJ84 | -0,001364 | 0,002321 | -0,000957 |
| Q9VGQ4 | -0,002577 | 0,004305 | -0,001727 |
| Q7KE33 | -0,008971 | 0,018651 | -0,009679 |
| E1JHS6 | -0,001604 | 0,004198 | -0,002593 |
| Q9VHJ0 | -0,007066 | 0,013451 | -0,006386 |
| Q8MSK0 | -0,027583 | 0,017123 | 0,010460 |
| B4ZJA2 | -0,009680 | 0,011453 | -0,001773 |
| Q6IGA5 | -0,000006 | 0,008270 | -0,008264 |
| Q8IQS7 | -0,002558 | 0,011984 | -0,009425 |
| C4NAP3 | -0,000686 | 0,000184 | 0,000503 |
| D1Z365 | -0,000640 | 0,002658 | -0,002018 |
| Q9VXU9 | -0,005530 | 0,008667 | -0,003137 |
| F0JAT6 | -0,010382 | 0,008451 | 0,001932 |
| Q9VWV6 | -0,007110 | 0,009701 | -0,002591 |
| Q6GUX9 | -0,000253 | 0,004972 | -0,004719 |
| Q9VJN9 | -0,000150 | 0,006757 | -0,006607 |
| Q3HKQ1 | -0,001726 | 0,001117 | 0,000609 |
| Q8MS62 | -0,005111 | 0,005528 | -0,000417 |
| B4ZJA4 | 0,024912 | 0,030855 | -0,055767 |
| Q9W2S7 | -0,007119 | 0,010506 | -0,003387 |
| Q9V3J3 | 0,000027 | 0,000033 | -0,000059 |

**Table S10. Bayesian posterior estimates of temperature effects on SFP production and transfer (short-term exposure / experiment 1 under no-SCR).** Values represent loge fold-change estimates of protein abundance induced by low (20ºC) and hot (28ºC) temperatures relative to the 24ºC baseline. For each temperature contrast, the posterior mean and 95% Credible Intervals (CrIs) are reported for A) Production and B) Transfer. $P(\beta > 0)$ denotes the posterior probability that the specific temperature contrast has a positive directional effect compared to the baseline. Estimates highlighted in bold indicate a highly probable, robust difference where the 95% CrI strictly excludes zero. Italicized values denote substantial directional evidence, where $P(\beta > 0) > 0.90$ indicates a probable increase, and $P(\beta > 0) < 0.10$ indicates a probable decrease in SFP abundance.

A.

| **Protein** | **20°C mean [95% CrI]** | **P(β > 0)** | **28°C mean [95% CrI]** | **P(β > 0)** |
|---|---|---|---|---|
| Sems | **1.43 [0.47, 2.26]** | *0.994* | **0.97 [0.01, 1.81]** | *0.976* |
| Lectin 46Cb | **1.44 [0.08, 2.61]** | *0.980* | 0.69 [−0.64, 1.87] | 0.887 |
| CG9997 | 1.33 [−0.38, 2.80] | *0.943* | 0.60 [−1.05, 2.12] | 0.796 |
| CG17575 | 1.11 [−0.74, 2.77] | *0.901* | 1.16 [−0.69, 2.74] | *0.912* |
| antr | 0.85 [−0.60, 2.17] | 0.898 | 0.60 [−0.83, 1.96] | 0.828 |
| Ovulin | 1.00 [−0.93, 2.76] | 0.862 | 0.32 [−1.53, 2.13] | 0.650 |
| Lectin 46Ca | 0.92 [−0.97, 2.59] | 0.858 | 1.05 [−0.82, 2.75] | 0.882 |
| aqrs | 0.48 [−0.74, 1.67] | 0.805 | 0.01 [−1.19, 1.23] | 0.511 |
| intr | 0.46 [−0.79, 1.63] | 0.800 | 0.19 [−1.09, 1.35] | 0.651 |
| Semp1 | −0.07 [−2.19, 2.06] | 0.480 | 1.04 [−1.18, 3.07] | 0.841 |
| SP | −0.14 [−2.55, 2.38] | 0.447 | −1.59 [−4.03, 1.06] | *0.106* |

B.

| **Protein** | **20°C mean [95% CrI]** | **P(β > 0)** | **28°C mean [95% CrI]** | **P(β > 0)** |
|---|---|---|---|---|
| CG9997 | **1.10 [0.27, 1.83]** | *0.992* | 0.18 [−0.66, 0.94] | 0.709 |
| Lectin 46 Cb | **1.00 [0.11, 1.81]** | *0.984* | 0.08 [−0.77, 0.90] | 0.589 |
| aqrs | 0.57 [−0.05, 1.17] | *0.968* | 0.02 [−0.61, 0.60] | 0.531 |
| Ovuline | 0.65 [−0.06, 1.34] | *0.966* | 0.28 [−0.42, 0.93] | 0.829 |
| intr | 0.52 [−0.14, 1.14] | *0.948* | -0.24 [−0.88, 0.41] | 0.208 |
| Sems | 0.92 [−0.25, 2.01] | *0.947* | -0.08 [−1.26, 1.07] | 0.434 |
| antr | 0.43 [−0.19, 1.07] | *0.923* | 0.05 [−0.58, 0.66] | 0.585 |
| Lectin 46Ca | 0.49 [−1.19, 2.08] | 0.745 | 0.60 [−1.07, 2.19] | 0.779 |
| CG17575 | 0.64 [−1.14, 2.23] | 0.791 | 0.55 [−1.22, 2.21] | 0.757 |
| SP | 0.00 [−1.16, 1.16] | 0.512 | 0.52 [−0.68, 1.61] | 0.834 |
| Semp1 | 0.00 [−1.64, 1.54] | 0.497 | 0.20 [−1.48, 1.77] | 0.618 |

**Table S11. Bayesian posterior estimates of temperature effects on SFP production and transfer (short-term exposure / experiment 1 under High-SCR).** Values represent loge fold-change estimates of protein abundance induced by low (20ºC) and hot (28ºC) temperatures relative to the 24ºC baseline. For each temperature contrast, the posterior mean and 95% Credible Intervals (CrIs) are reported for A) Production and B) Transfer. $P(\beta > 0)$ denotes the posterior probability that the specific temperature contrast has a positive directional effect compared to the baseline. Estimates highlighted in bold indicate a highly probable, robust difference where the 95% CrI strictly excludes zero. Italicized values denote substantial directional evidence, where $P(\beta > 0) > 0.90$ indicates a probable increase, and $P(\beta > 0) < 0.10$ indicates a probable decrease in SFP abundance.

A.

| **Protein** | **20°C mean [95% CrI]** | **P(β > 0)** | **28°C mean [95% CrI]** | **P(β > 0)** |
|---|---|---|---|---|
| Lectin 46Cb | 0.88 [−0.19, 2.11] | *0.952* | 0.60 [−0.55, 1.70] | 0.875 |
| Lectin 46Ca | −0.45 [−1.90, 1.07] | 0.254 | 0.92 [−0.54, 2.31] | *0.902* |
| SP | −1.89 [−3.91, 0.25] | *0.040* | 0.50 [−1.94, 2.61] | 0.698 |
| CG17575 | 0.20 [−1.87, 2.49] | 0.563 | 0.36 [−1.55, 2.31] | 0.638 |
| Semp1 | −0.07 [−1.54, 1.56] | 0.463 | 0.21 [−1.34, 2.03] | 0.596 |
| Ovulin | −0.72 [−2.16, 0.85] | 0.166 | −0.12 [−1.78, 1.42] | 0.450 |
| CG9997 | −0.16 [−2.11, 1.69] | 0.433 | −0.43 [−2.76, 1.38] | 0.323 |
| aqrs | −0.34 [−2.04, 1.36] | 0.331 | −0.28 [−1.90, 1.39] | 0.366 |
| antr | −0.21 [−1.72, 1.38] | 0.382 | −0.53 [−2.07, 1.01] | 0.227 |
| intr | −0.17 [−0.86, 0.56] | 0.271 | −0.09 [−0.76, 0.67] | 0.353 |
| Sems | −0.41 [−1.80, 1.01] | 0.274 | −0.18 [−1.64, 1.16] | 0.400 |

B.

| **Protein** | **20°C mean [95% CrI]** | **P(β > 0)** | **28°C mean [95% CrI]** | **P(β > 0)** |
|---|---|---|---|---|
| Lectin 46Ca | −0.15 [−1.24, 0.90] | 0.377 | **1.01 [−0.07, 2.08]** | *0.968* |
| Lectin 46Cb | 0.53 [−0.43, 1.43] | 0.886 | 0.37 [−0.57, 1.29] | 0.811 |
| Semp1 | 0.14 [−1.48, 1.70] | 0.576 | 0.05 [−1.49, 1.65] | 0.523 |
| SP | −1.02 [−3.56, 1.62] | 0.204 | 0.21 [−2.41, 2.84] | 0.562 |
| CG17575 | −0.56 [−1.93, 0.87] | 0.200 | −0.04 [−1.40, 1.38] | 0.465 |
| aqrs | −0.36 [−1.70, 1.11] | 0.271 | −0.15 [−1.53, 1.23] | 0.407 |
| CG9997 | −0.34 [−1.97, 1.29] | 0.331 | −0.16 [−1.94, 1.50] | 0.435 |
| Ovulin | −0.22 [−1.57, 1.18] | 0.359 | −0.11 [−1.40, 1.27] | 0.430 |
| antr | −0.25 [−0.90, 0.39] | 0.192 | −0.21 [−0.86, 0.43] | 0.246 |
| intr | −0.28 [−0.89, 0.33] | 0.167 | −0.14 [−0.74, 0.48] | 0.296 |
| Sems | −0.25 [−1.04, 0.54] | 0.237 | −0.05 [−0.82, 0.75] | 0.440 |

**Table S12. Bayesian posterior estimates of temperature effects on SFP production and transfer (short-term exposure / experiment 2 under No-SCR).** Values represent loge fold-change estimates of protein abundance induced by low (20ºC) and hot (28ºC) temperatures relative to the 24ºC baseline. For each temperature contrast, the posterior mean and 95% Credible Intervals (CrIs) are reported for A) Production and B) Transfer. P(β > 0) denotes the posterior probability that the specific temperature contrast has a positive directional effect compared to the baseline. Estimates highlighted in bold indicate a highly probable, robust difference where the 95% CrI strictly excludes zero. Italicized values denote substantial directional evidence, where P(β > 0) > 0.90 indicates a probable increase, and P(β > 0) < 0.10 indicates a probable decrease in SFP abundance.

A.

| **Protein** | **20°C mean [95% CrI]** | **P(β > 0)** | **28°C mean [95% CrI]** | **P(β > 0)** |
|---|---|---|---|---|
| intr | **−0.80 [−1.35, −0.21]** | *0.010* | −0.23 [−0.77, 0.34] | 0.181 |
| aqrs | −0.88 [−1.86, 0.16] | *0.040* | −0.18 [−1.13, 0.78] | 0.329 |
| CG9997 | −1.24 [−2.64, 0.34] | 0.055 | −0.71 [−2.13, 0.79] | 0.152 |
| CG17575 | −1.32 [−3.03, 0.48] | 0.069 | 0.68 [−1.08, 2.40] | 0.795 |
| antr | −0.61 [−1.34, 0.21] | *0.058* | −0.39 [−1.12, 0.39] | 0.139 |
| Ovulin | −0.94 [−2.56, 0.80] | 0.121 | −0.62 [−2.18, 1.05] | 0.214 |
| SP | −0.80 [−3.59, 2.12] | 0.289 | −0.20 [−3.01, 2.74] | 0.435 |
| Sems | −0.66 [−1.93, 0.66] | *0.149* | −0.36 [−1.66, 1.01] | 0.269 |
| Semp1 | −0.57 [−2.86, 1.78] | 0.301 | 0.50 [−1.87, 2.82] | 0.672 |
| Lectin 46Ca | −0.51 [−2.67, 1.76] | 0.313 | −0.25 [−2.45, 2.00] | 0.406 |
| Lectin 46Cb | −0.43 [−1.88, 1.04] | 0.259 | 0.45 [−1.04, 1.90] | 0.752 |

B.

| **Protein** | **20°C mean [95% CrI]** | **P(β > 0)** | **28°C mean [95% CrI]** | **P(β > 0)** |
|---|---|---|---|---|
| CG17575 | −1.36 [−2.88, 0.27] | *0.045* | −0.05 [−1.57, 1.54] | 0.466 |
| Ovulin | −0.82 [−1.78, 0.23] | 0.054 | −0.85 [−1.85, 0.18] | *0.046* |
| CG9997 | −0.82 [−1.88, 0.37] | *0.073* | −1.01 [−2.10, 0.18] | *0.043* |
| Lectin 46Ca | −1.06 [−2.57, 0.63] | 0.089 | −0.90 [−2.41, 0.72] | 0.120 |
| Lectin 46Cb | −0.56 [−1.38, 0.28] | *0.084* | 0.21 [−0.59, 1.03] | 0.716 |
| Sems | −0.68 [−1.66, 0.35] | *0.079* | −0.31 [−1.27, 0.67] | 0.242 |
| SP | −0.79 [−2.19, 0.78] | 0.132 | −0.75 [−2.20, 0.76] | 0.140 |
| Semp1 | −0.83 [−3.09, 1.52] | 0.228 | 0.28 [−2.02, 2.63] | 0.602 |
| antr | −0.12 [−0.80, 0.62] | 0.338 | −0.51 [−1.19, 0.21] | *0.063* |
| aqrs | −0.14 [−1.24, 1.02] | 0.383 | −0.47 [−1.54, 0.66] | 0.166 |
| intr | −0.28 [−1.03, 0.48] | 0.205 | −0.35 [−1.10, 0.38] | 0.147 |

**Table S13. Bayesian posterior estimates of temperature effects on SFP production and transfer (short-term exposure / experiment 2 under High-SCR).** Values represent $\log_e$ fold-change estimates of protein abundance induced by low (20ºC) and hot (28ºC) temperatures relative to the 24ºC baseline. For each temperature contrast, the posterior mean and 95% Credible Intervals (CrIs) are reported for A) Production and B) Transfer. $P(\beta > 0)$ denotes the posterior probability that the specific temperature contrast has a positive directional effect compared to the baseline. Estimates highlighted in bold indicate a highly probable, robust difference where the 95% CrI excludes zero. Italicized values denote substantial directional evidence, where $P(\beta > 0) > 0.90$ indicates a probable increase, and $P(\beta > 0) < 0.10$ indicates a probable decrease in SFP abundance.

A.

| **Protein** | **20°C mean [95% CrI]** | **P(β > 0)** | **28°C mean [95% CrI]** | **P(β > 0)** |
|---|---|---|---|---|
| CG17575 | −1.81 [−3.98, 0.33] | *0.044* | −1.13 [−3.38, 1.03] | 0.133 |
| SP | −2.52 [−7.65, 2.47] | 0.144 | −2.23 [−7.27, 2.71] | 0.172 |
| Semp1 | −1.20 [−3.65, 1.36] | 0.143 | −1.28 [−3.75, 1.19] | 0.123 |
| Ovulin | −1.19 [−3.02, 0.58] | *0.077* | −0.92 [−2.75, 0.80] | 0.126 |
| antr | −0.79 [−2.11, 0.44] | *0.081* | −0.93 [−2.19, 0.26] | *0.056* |
| Lectin 46Ca | −0.56 [−2.99, 1.84] | 0.304 | −1.07 [−3.47, 1.29] | 0.154 |
| Lectin 46Cb | −0.62 [−2.48, 1.17] | 0.225 | −0.98 [−2.76, 0.81] | 0.116 |
| aqrs | −0.53 [−2.34, 1.30] | 0.250 | −0.06 [−1.85, 1.69] | 0.485 |
| intr | −0.47 [−3.05, 2.16] | 0.340 | −0.37 [−2.94, 2.19] | 0.369 |
| CG9997 | −0.69 [−3.52, 2.00] | 0.288 | −0.58 [−3.47, 2.09] | 0.318 |
| Sems | −0.55 [−2.63, 1.46] | 0.276 | −0.69 [−2.71, 1.39] | 0.224 |

B.

| **Protein** | **20°C mean [95% CrI]** | **P(β > 0)** | **28°C mean [95% CrI]** | **P(β > 0)** |
|---|---|---|---|---|
| CG17575 | **−2.38 [−3.16, −1.58]** | *~0.000* | **−2.22 [−2.99, −1.43]** | *~0.000* |
| Lectin 46Ca | **−1.39 [−2.13, −0.62]** | *0.001* | **−1.19 [−1.94, −0.44]** | *0.004* |
| Lectin 46Cb | **−0.99 [−2.04, 0.04]** | *0.030* | **−1.20 [−2.23, −0.18]** | *0.016* |
| Sems | **−1.16 [−2.32, 0.04]** | *0.028* | **−1.16 [−2.28, 0.02]** | *0.027* |
| Semp1 | −1.35 [−3.54, 0.83] | *0.096* | −1.73 [−3.89, 0.36] | *0.047* |
| antr | −0.44 [−1.31, 0.42] | 0.128 | −0.37 [−1.21, 0.52] | 0.170 |
| SP | −1.01 [−3.74, 1.79] | 0.207 | −0.45 [−3.20, 2.34] | 0.354 |
| Ovulin | −0.66 [−2.14, 0.76] | 0.158 | −0.42 [−1.82, 0.93] | 0.244 |
| aqrs | −0.50 [−1.70, 0.70] | 0.175 | −0.12 [−1.35, 1.11] | 0.407 |
| intr | −0.43 [−1.54, 0.70] | 0.189 | −0.21 [−1.30, 0.94] | 0.333 |
| CG9997 | −0.44 [−1.81, 0.94] | 0.231 | −0.29 [−1.58, 1.06] | 0.308 |

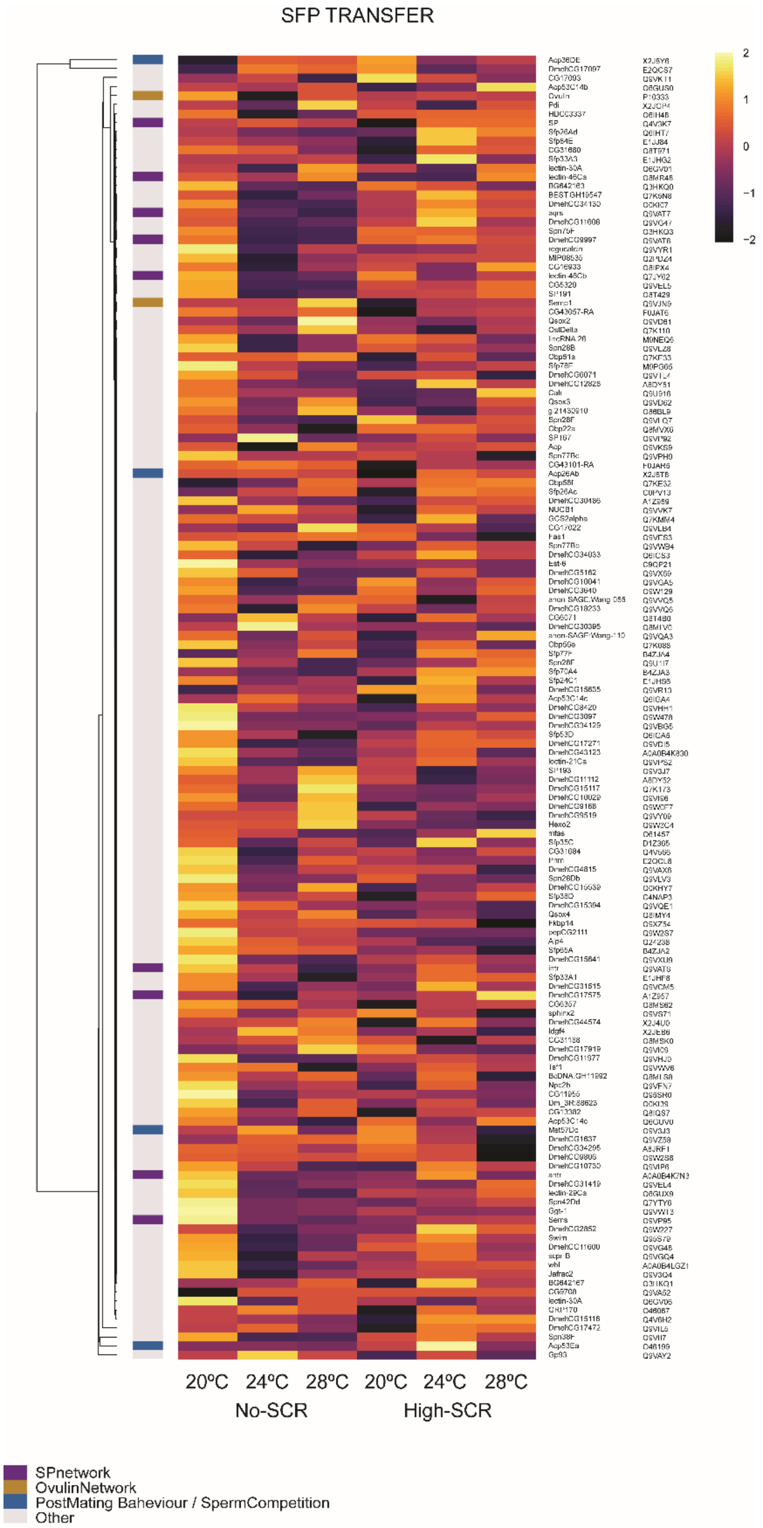


**Figure S3.** Heatmaps showing absolute abundance–based transfer estimates for the 145 SFPs detected in accessory gland samples (short-term exposure/experiment 1). Each cell gives the across-biological replicate mean for that protein, at each temperature treatment. Row annotations indicate functional information relating to protein functions as part of the sex peptide or ovulin networks, or other known roles in sperm competition.

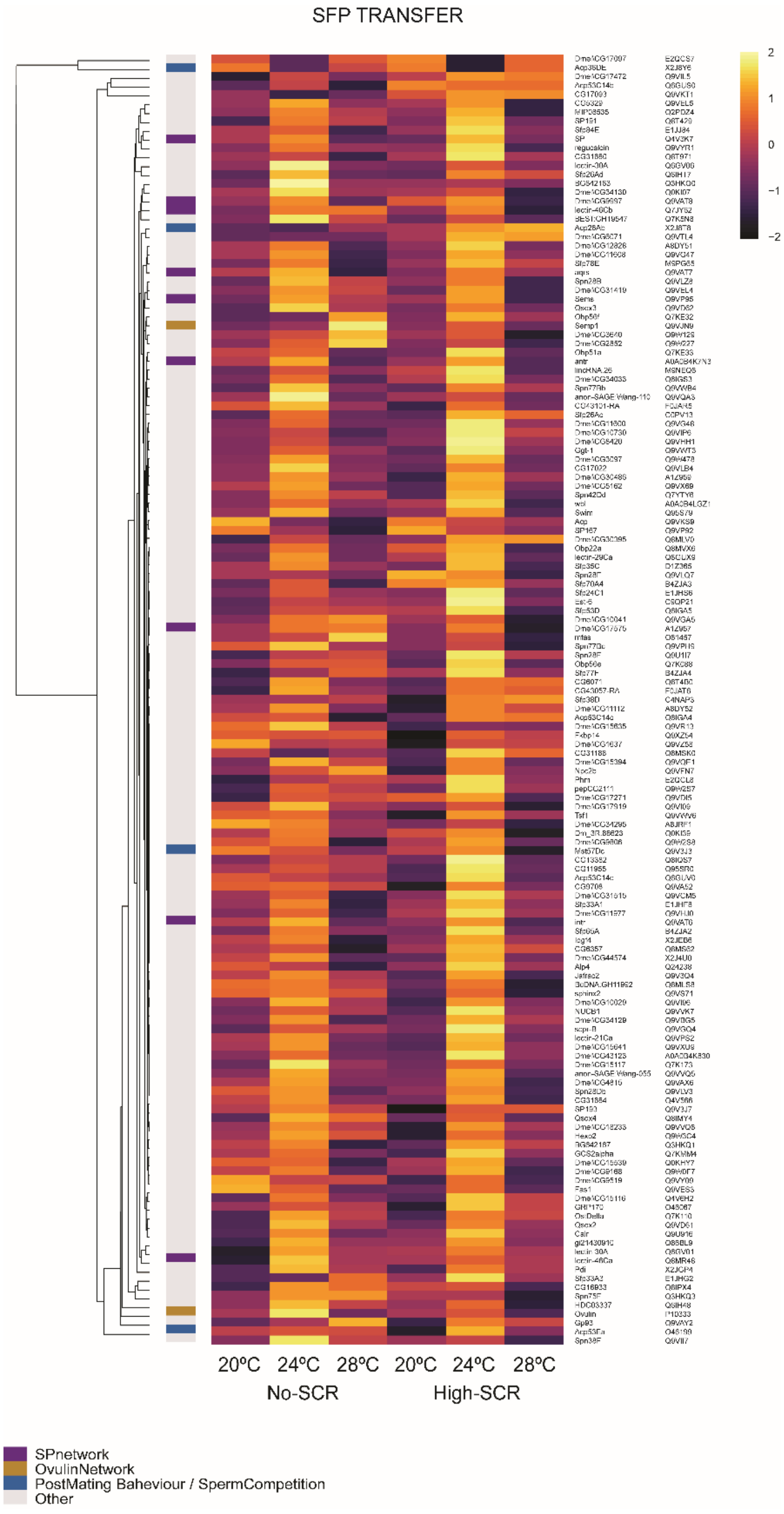


**Figure S4.** Heatmaps showing absolute abundance–based transfer estimates for the 145 SFPs detected in accessory gland samples (long-term exposure/experiment 2). Each cell gives the across-biological replicate mean for that protein, at each temperature treatment. Row annotations indicate functional information relating to protein functions as part of the sex peptide or ovulin networks, or other known roles in sperm competition.

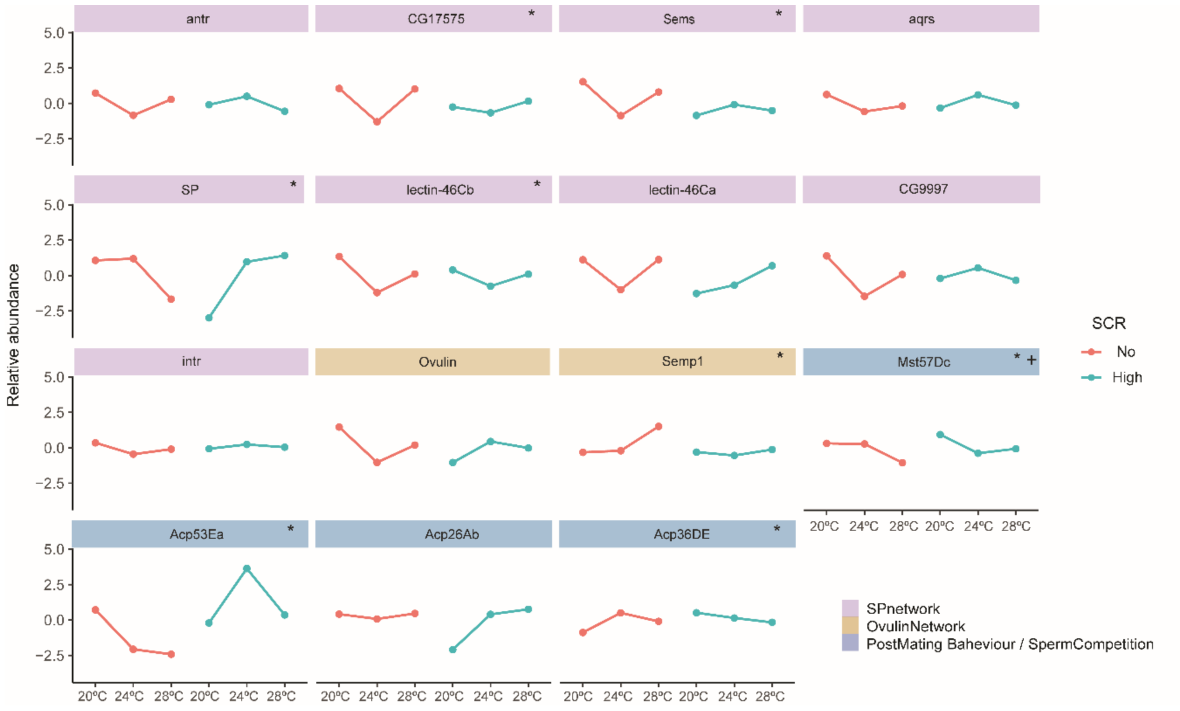


**Figure S5.** Relative abundance production profiles of SFPs with key function on female post-mating responses after 48hrs of treatment duration (experiment 1). Each point represents an average across the 3 replicates in relation to each temperature and sperm competition risk. Abundance values were normalized by mean-centering and averaged across replicates. *Proteins selected as important having the strongest contribution to temperature responses under no-SCR. + Proteins selected as important under high-SCR.

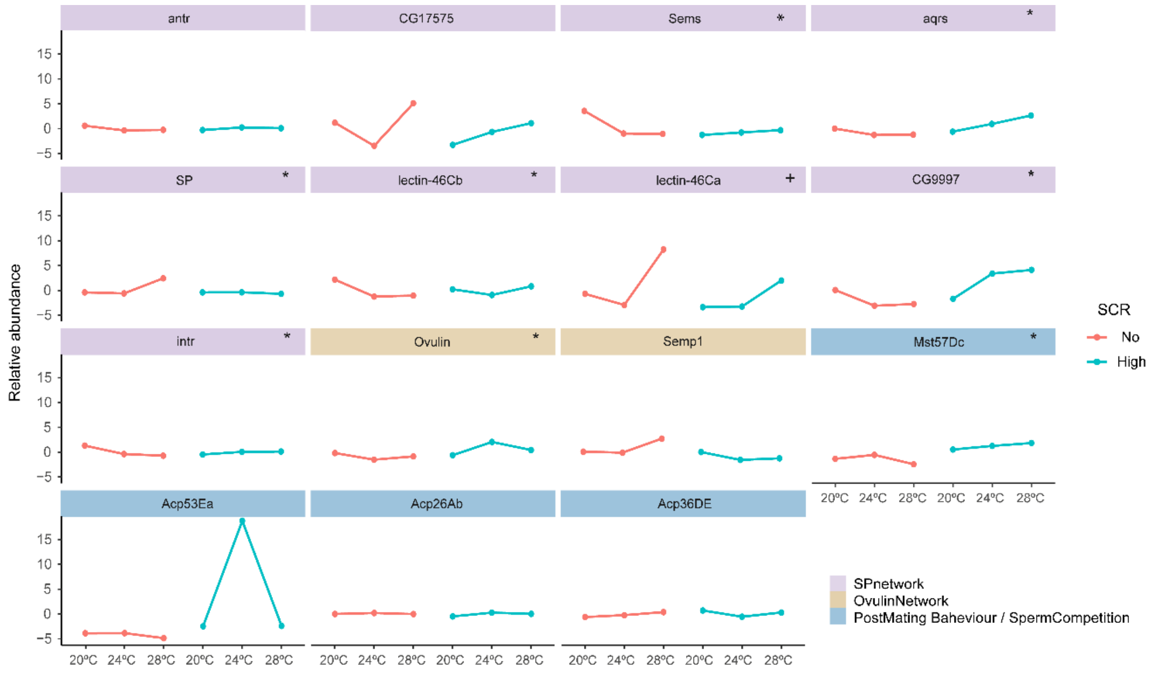


**Figure S6.** Relative abundance transfer profiles of SFPs with key function on female post-mating responses after 48hrs of treatment duration (experiment 1). Each point represents an average across the 3 replicates in relation to each temperature and sperm competition risk. Abundance values were normalized by mean-centering and averaged across replicates. *Proteins selected as important having the strongest contribution to temperature responses under no-SCR. + Proteins selected as important under high-SCR.

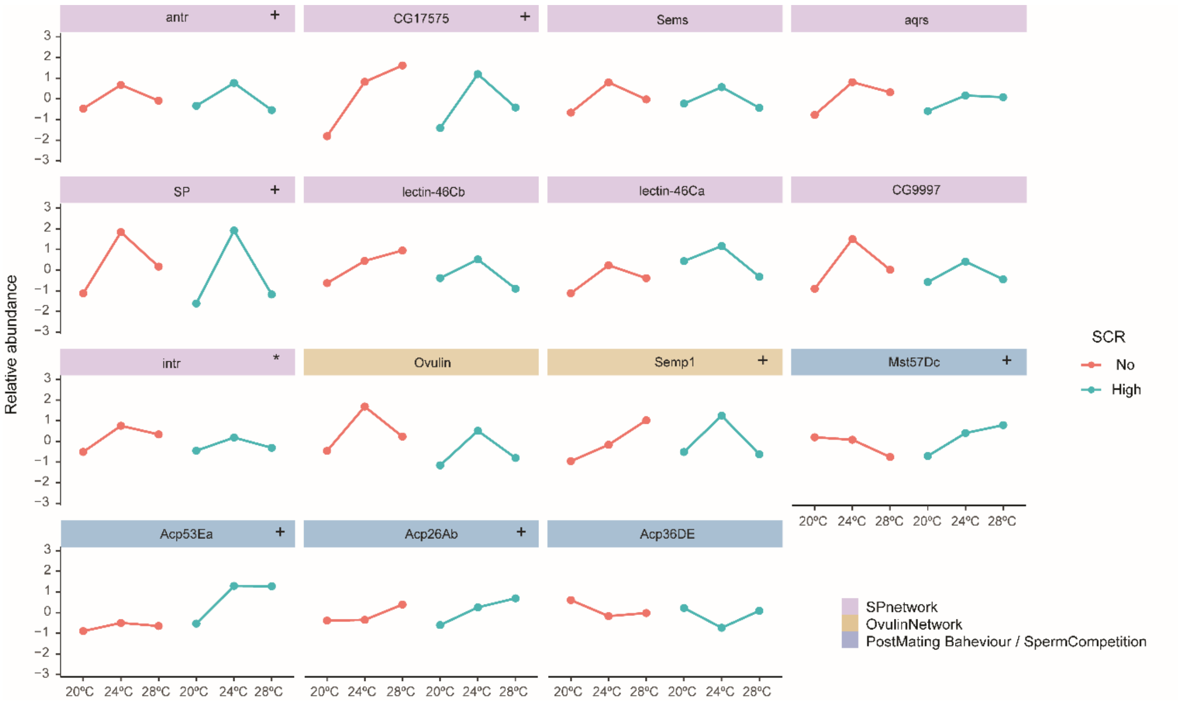


**Figure S7.** Relative abundance production profiles of SFPs with key function on female post-mating responses after 13 days of treatment duration (experiment 2). Each point represents an average across the 3 replicates in relation to each temperature and sperm competition risk. Abundance values were normalized by mean-centering and averaged across replicates. *Proteins selected as important having the strongest contribution to temperature responses under no-SCR. + Proteins selected as important under high-SCR.

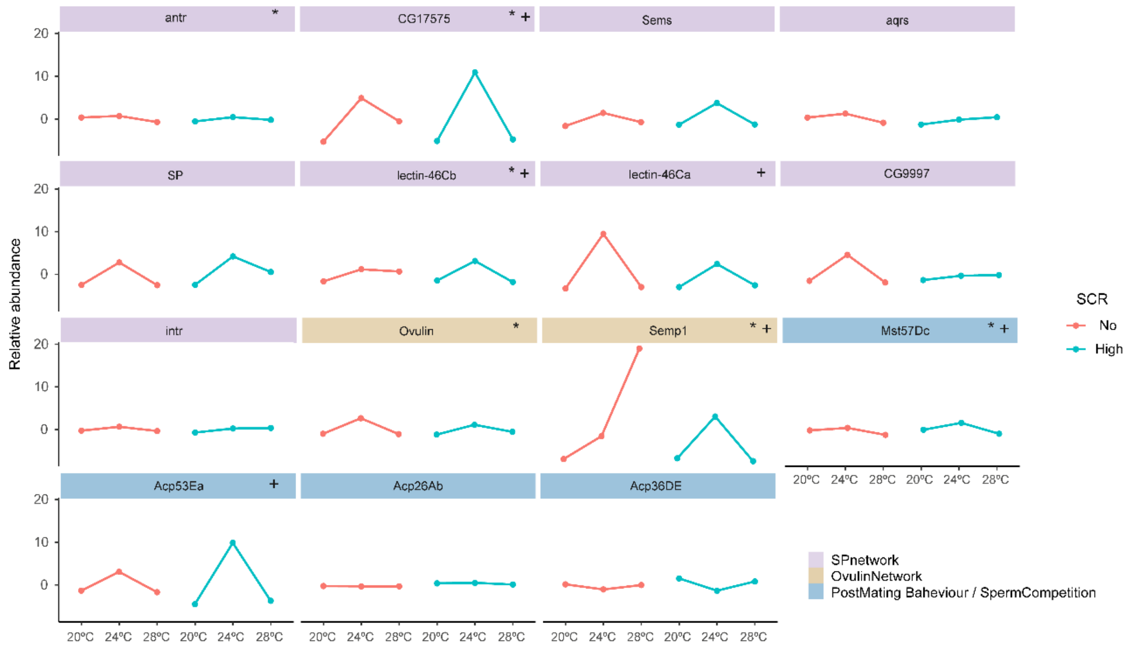


**Figure S8.** Relative abundance transfer profiles of SFPs with key function on female post-mating responses after 13 days of treatment duration (experiment 2). Each point represents an average across the 3 replicates in relation to each temperature and sperm competition risk. Abundance values were normalized by mean-centering and averaged across replicates. *Proteins selected as important having the strongest contribution to temperature responses under no-SCR. + Proteins selected as important under high-SCR.

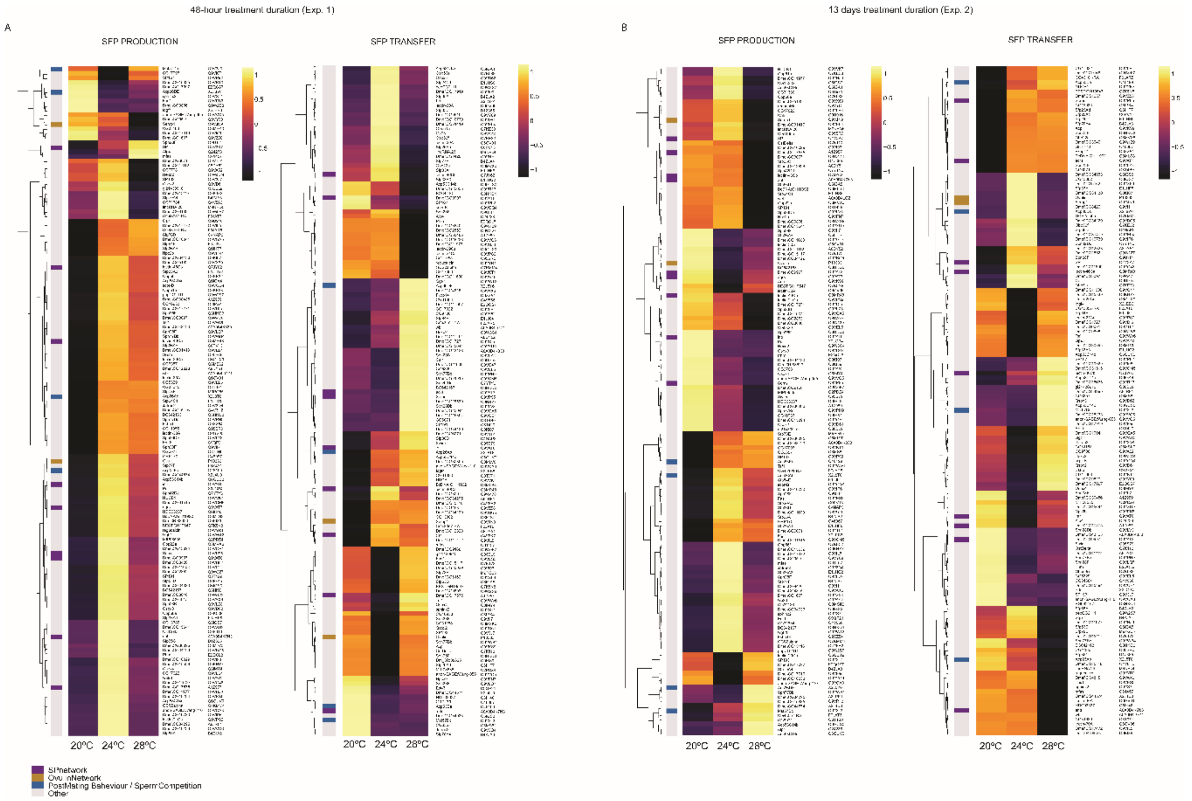


**Figure S9 | Comparison of SFPs production and transfer under high-SCR vs. no-SCR across temperature treatments.** A) Short-term exposure / experiment 1. B) Long-term exposure / experiment 2. Each cell shows the mean ratio of protein absolute abundance under high-SCR relative to no-SCR across biological replicates for each temperature. Values closer to 1 indicate higher production/transfer under high-SCR conditions. Row annotations indicate functional information relating to protein functions as part of the sex peptide or ovulin networks, or other known roles in sperm competition.

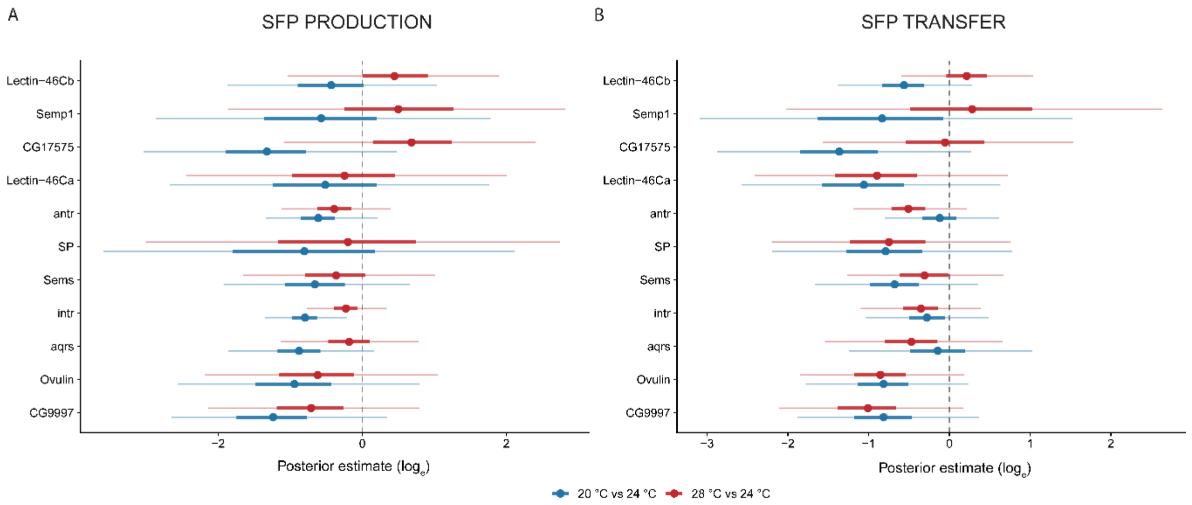


**Figure S10 | Bayesian posterior estimates of temperature effect on production and transfer of SFPs involved in Ovulin and SP networks (long-term exposure / experiment 2 under no-SCR)**. $\log_e$ fold-change estimates of protein abundance induced by low (20ºC) and hot (28ºC) temperatures relative to the 24ºC baseline. A) Posterior estimates for SFP production. B) Posterior estimates for SFP transfer. Blue intervals represent the 20ºC vs 24ºC contrast, while red intervals represent the 28ºC vs 24ºC contrast. Central point on each segment denotes the median posterior estimate for a specific protein. Thick and thin horizontal lines represent the 50% and 95% Credible Intervals (CIs), respectively. Vertical dashed line at zero represents no change in abundance compared to the 24ºC baseline. Estimates where the 95% CI strictly excludes zero indicate a highly probable (>95%), robust difference induced by the specific temperature treatment.

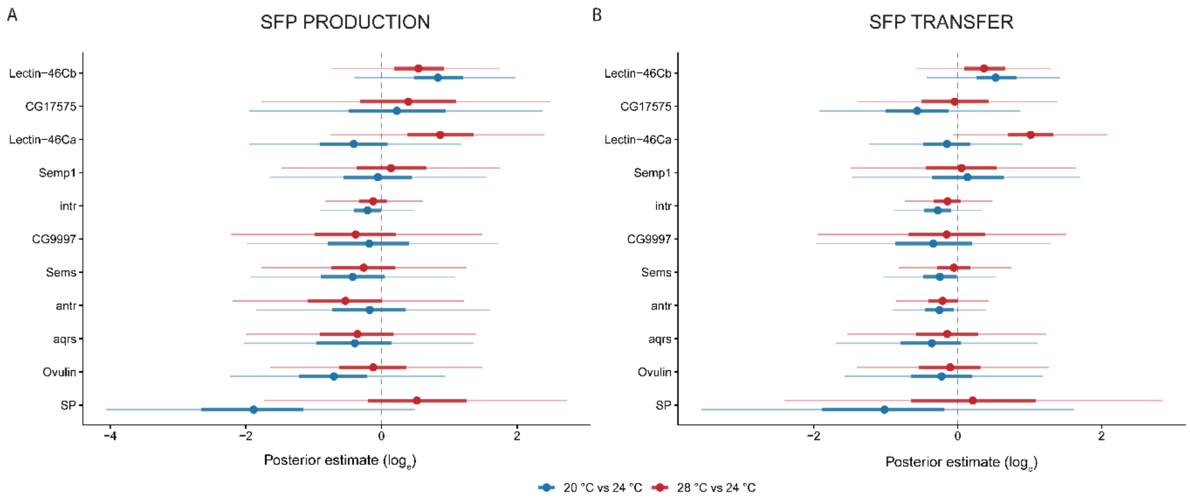


**Figure S11 | Bayesian posterior estimates of temperature effect on production and transfer of SFPs involved in Ovulin and SP networks (short-term exposure / experiment 1 under high-SCR)**. log$_e$ fold-change estimates of protein abundance induced by low (20ºC) and hot (28ºC) temperatures relative to the 24ºC baseline. A) Posterior estimates for SFP production. B) Posterior estimates for SFP transfer. Blue intervals represent the 20ºC vs 24ºC contrast, while red intervals represent the 28ºC vs 24ºC contrast. Central point on each segment denotes the median posterior estimate for a specific protein. Thick and thin horizontal lines represent the 50% and 95% Credible Intervals (CIs), respectively. Vertical dashed line at zero represents no change in abundance compared to the 24ºC baseline. Estimates where the 95% CI strictly excludes zero indicate a highly probable (>95%), robust difference induced by the specific temperature treatment.
